# Immunological and pathological outcomes of SARS-CoV-2 challenge after formalin-inactivated vaccine immunisation of ferrets and rhesus macaques

**DOI:** 10.1101/2020.12.21.423746

**Authors:** Kevin R. Bewley, Karen Gooch, Kelly M. Thomas, Stephanie Longet, Nathan Wiblin, Laura Hunter, Kin Chan, Phillip Brown, Rebecca A. Russell, Catherine Ho, Gillian Slack, Holly E. Humphries, Leonie Alden, Lauren Allen, Marilyn Aram, Natalie Baker, Emily Brunt, Rebecca Cobb, Susan Fotheringham, Debbie Harris, Chelsea Kennard, Stephanie Leung, Kathryn Ryan, Howard Tolley, Nadina Wand, Andrew White, Laura Sibley, Charlotte Sarfas, Geoff Pearson, Emma Rayner, Xiaochao Xue, Teresa Lambe, Sue Charlton, Sarah Gilbert, Quentin J. Sattentau, Fergus Gleeson, Yper Hall, Simon Funnell, Sally Sharpe, Francisco J. Salguero, Andrew Gorringe, Miles Carroll

## Abstract

There is an urgent requirement for safe and effective vaccines to prevent novel coronavirus disease (COVID-19) caused by SARS-CoV-2. A concern for the development of new viral vaccines is the potential to induce vaccine-enhanced disease (VED). This was reported in several preclinical studies with both SARS-CoV-1 and MERS vaccines but has not been reported with SARS-CoV-2 vaccines. We have used ferret and rhesus macaques challenged with SARS-CoV-2 to assess the potential for VED in animals vaccinated with formaldehyde-inactivated SARS-CoV-2 (FIV) formulated with Alhydrogel, compared to a negative control vaccine in ferrets or unvaccinated macaques. We showed no evidence of enhanced disease in ferrets or rhesus macaques given FIV except for mild transient enhanced disease seen at seven days post infection in ferrets. This increased lung pathology was observed early in the infection (day 7) but was resolved by day 15. We also demonstrate that formaldehyde treatment of SARS-CoV-2 reduces exposure of the spike receptor binding domain providing a mechanistic explanation for suboptimal immunity.

## Introduction

Novel coronavirus disease (COVID-19) caused by SARS-CoV-2 is a global pandemic with a cumulative total of over 63 million cases and 1.4 million deaths reported as of 2^nd^ December 2020^1^. Consequently, there is an urgent requirement to develop safe and effective vaccines to prevent COVID-19^2^. Currently 52 vaccine candidates are in clinical evaluation (11 at Phase 3) with 162 listed as in pre-clinical evaluation (WHO draft landscape of COVID-19 vaccines – 8 December 2020). The leading vaccine candidates in Phase 3 studies include a non-replicating viral vector, three inactivated virus vaccines and two vaccines based on mRNA technology^3^. Promising clinical results from phase 2/3 studies are available for mRNA-based vaccines^4 5^, the adenovirus vaccine (ChAdOx1nCoV-19/AZD1222)^6^, which expresses a codon-optimised full-length spike protein (S) and whole virus vaccines grown in Vero cells and inactivated with β-propiolactone^7 8 9^. These vaccines have also been evaluated for protection in non-human primates following challenge with SARS-CoV-2. The virus induces only mild to moderate disease in macaques but these vaccines reduce viral loads and pathology in the upper and lower respiratory tracts to varying degrees^10 8 7 11^.

A concern for the development of new viral vaccines is the potential to induce vaccine enhanced disease^12^ (VED) which has been associated with prior pre-clinical studies of both SARS and MERS vaccines. The most studied mechanism of VED is antibody-dependent enhancement (ADE) of disease, reviewed recently by Arvin *et al*.^13^. It has been suggested that ADE could be a consequence of low affinity antibodies that bind to viral proteins but have limited neutralising activity^14^. The vaccine enhancement of disease by ADE mechanisms was described in children given formaldehyde-inactivated respiratory syncytial virus (RSV) vaccines in the 1960s^15^, measles vaccines^16^ and in dengue haemorrhagic fever due to secondary infection with a heterologous dengue serotype^17^.

There is limited evidence of ADE with SARS-CoV-1 vaccines in animal models and whilst it has not been reported in the majority of vaccine studies, a study that used formalin or ultraviolet-inactivated SARS-CoV-1 observed that older mice developed pulmonary pathology with an eosinophil infiltrate^18^. A further study demonstrated protection in mice following immunisation with formalin or ultraviolet light-inactivated SARS-CoV-1, but animals developed eosinophilic pulmonary infiltrates^19^. A modified vaccinia virus Ankara expressing S protein (MVA-S) was not protective in ferrets challenged with SARS-CoV-1, but liver inflammation was noted^20^. Formalin-inactivated SARS-CoV-1 vaccines were protective in rhesus macaques^21^, but also promoted lymphocytic infiltrates and alveolar oedema with fibrin deposition after challenge^22^. Likewise, MVA expressing S protein showed protection in one study^23^, but greater occurrence of diffuse alveolar damage than seen in control animals following challenge. Fortuitously, VED has not been reported in the numerous SARS-CoV-2 efficacy animal challenge vaccine studies published to date. However, these studies were primarily designed to assess efficacy and not VED.

We have developed a ferret intranasal SARS-CoV-2 infection model where viral shedding and mild lung pathology is observed and re-challenged animals are fully protected^24^. We have also evaluated both rhesus and cynomolgus macaques for their susceptibility to SARS-CoV-2 challenge and showed the development of pulmonary lesions in both species which are equivalent to those seen in mild clinical cases in humans^25^. In order to interrogate the potential for VED in ferrets and rhesus macaques to support future safety studies on novel COVID-19 vaccines, we prepared a formaldehyde-inactivated SARS-CoV-2 vaccine (FIV), formulated in Alhydrogel. In this study design we aim to induce a suboptimal immune response which may promote VED, immunised animals are challenged with SARS-CoV-2 14 days after vaccination. Clinical signs, viral shedding and pathology are monitored following challenge, and immune responses characterised before and after infection. No enhanced pathology is observed in either species except for transient enhanced pathology at 7 days post infection in ferrets and we present a possible mechanism for suboptimal immunity induced by formaldehyde-inactivated SARS-CoV-2 spike.

## Materials & Methods

### Viruses and cells

SARS-CoV-2 Victoria/01/2020^26^ was provided by The Doherty Institute, Melbourne, Australia at P1 and passaged twice in Vero/hSLAM cells [ECACC 04091501]. Briefly, confluent monolayers of hSLAM cells were infected at a multiplicity of infection (MOI) of approximately 0.0005 for 60 min in medium (see below) containing no FBS at 37°C. The flasks were then filled with media supplemented with 4% heat-inactivated FBS. Virus was harvested at 72 h post-infection by removal of any remaining attached cells with sterile 5 mm borosilicate glass beads, clarification of the cell/media supernatant by centrifugation at 1000 x g for 10 min, followed by dispensing and storage at ≥-65°C. Whole genome sequencing was performed, on the challenge isolate, using both Nanopore and Illumina as described previously^24^. Virus titre was determined by plaque assay on Vero/E6 cells [ECACC 85020206]. Cell cultures were maintained at 37°C in MEM (Life Technologies, California, USA) supplemented with 10% foetal bovine serum (FBS, Sigma, Dorset, UK) and 25mM HEPES (Life Technologies). All Vero/hSLAM cell cultures were also supplemented with 0.4 μg/ml Geneticin (Gibco).

### Preparation of formalin-inactivated virus vaccine (FIV)

Centrifugal concentrators (VivaSpin20; 300kDa cut off) were sterilised with 20 mL of 70% ethanol for 10 min followed by a wash with 20mL of Dulbecco’s PBS (Gibco). To reduce the concentration of calf serum components in the material, concentrators were loaded with 120mL of SARS-CoV-2 at a titre of 8.45 x 10^6^ pfu/mL and centrifuged at 3000 x *g* for 60 – 80 min (until the retained volume was ≤ 2mL); the concentrators were re-filled with 20mL of DPBS and centrifugation repeated for a total of three washes. After the final wash, the material was pooled and made up to 30mL in sterile DPBS. Methanol-free formaldehyde solution at 36% (w/v) was added to a final formaldehyde concentration of 0.02% at room temperature for 72 h. The inactivated virus was subjected to a further three 20mL DPBS washes to remove the residual formaldehyde, made up to 20mL in DPBS and aliquoted and stored below −15°C. To confirm inactivation, virus was seeded onto Vero/hSLAM cells in three flasks (100μl/flask) which were serially passaged for a total of 27 days. Microscopic examination for signs of cytopathic effect (CPE) and RTqPCR were used to confirm no viable virus remained.

### Assessment of inactivated SARS-CoV-2

SDS PAGE: Samples were added to Laemmli buffer (Sigma, S3401) and heated at 90°C for 5 min and loaded onto a 10-well NuPAGE 4-12% Bis-Tris gel, 1.0mm (ThermoFisher). 5μL SeeBlue Plus2 (ThermoFisher) ladder was loaded as a marker and gels were stained with SimplyBlue SafeStain (ThermoFisher). Western Blot: Samples were processed as described for SDS PAGE and transferred to PVDF membrane with iBlot2 (ThermoFisher). After transfer, membranes were washed with tris-buffered saline 0.1% Tween20 (TBST) for 5 min at room temperature, followed by 1 h in blocking buffer (TBST, 5% skimmed milk powder). Membranes were washed three times for 5 min with TBST. MERS convalescent neutralising serum (NIBSC S3) was diluted 1:1000 in blocking buffer and incubated for 1 h at room temperature and then at 4°C overnight. Membranes were washed three times for 5 min with TBST and then incubated with either anti-human IgG-AP or anti-rabbit IgG-AP (1:5000 in blocking buffer) for 1 h with agitation. Membranes were washed three times as above and then developed with BCIP/NBT liquid substrate system (Sigma-Aldrich). The protein concentration of the FIV was determined using a BCA assay (Pierce #23227) (859μg/mL). Densitometry analysis (ImageQuant TL; GE Healthcare) of the Western blot revealed a band of approximately 180 kDa that was only present in wild-type virus and FIV preparations. The relative density of this band (20.7%) permitted estimation of the proportion of the FIV total protein that was Coronavirus-specific (178μg/mL).

### Transmission electron microscopy

Live virus was inactivated and fixed with final concentrations of 4%(w/v) formaldehyde and 2.5%(w/v) glutaraldehyde at ambient temperature for >16 h prior to processing. Inactivated virus was processed without any additional fixation steps. Samples (approximately 10μL) were placed directly on to electron microscopy grids (400 mesh copper grid, covered with a carbon reinforced plastic film). After 5 min adsorption the sample was removed, and the grids were negatively stained using 2% methylamine tungstate. The grids were examined using a CM100 transmission electron microscope (Philips/FEI/ThermoFisher Scientific) operated at 80kV.

### Animals. Ferrets

Ten healthy, female ferrets (*Mustela putorius furo*) aged 5-7 months were obtained from a UK Home Office accredited supplier (Highgate Farm, UK). The mean weight at the time of challenge was 1002g/ferret (range 871-1150g). Animals were housed as described previously^24^.

### Rhesus macaques

Twelve rhesus macaques of Indian origin (*Macaca mulatta*) were used in the study. Study groups comprised three males and three females and all were adults aged 2-4 years and weighing between 3.73 and 5.52 kg at the time of challenge. Animals were housed as described previously^25^. All experimental work was conducted under the authority of a UK Home Office approved project licence that had been subject to local ethical review at PHE Porton Down by the Animal Welfare and Ethical Review Body (AWERB).

### Vaccinations

Animals were randomly assigned to control (Ad-GFP for ferrets, no vaccine for rhesus macaques) and FIV-vaccinated groups. The weight distribution of the ferrets was tested to ensure there was no difference between groups (t-test, p> 0.05). An identifier chip (Bio-Thermo Identichip, Animalcare Ltd, UK) was inserted subcutaneously into the dorsal cervical region of each animal. Macaques were stratified for sex and into socially compatible cohorts and then randomly assigned into treatment groups. FIV was diluted in PBS to 133μg/mL Coronavirus-specific protein and mixed 1:1 in 2% Alhydrogel (Invivogen vac-alu-250) to give a final concentration of 66.7μg/mL in 1% Alhydrogel. Ferrets were immunised with a single intramuscular dose of 10μg of Coronavirus-specific protein in 150μL divided over two sites and macaques were immunised with 25 μg in 300 μL administered into the quadriceps femoris muscle of the right leg. Vaccination was 14 days before challenge. Control ferrets were immunised with a single intramuscular dose of 2.5 x 10^10^ virus particles of Ad-GFP^27^, a replication-deficient simian adenovirus vector containing an insert unrelated to Coronavirus (Green Fluorescent Protein, GFP), 28 days prior to challenge. Control macaques received no vaccine.

### SARS-CoV-2 challenge

Prior to challenge ferrets were sedated by intramuscular injection of ketamine/xylazine (17.9 mg/kg and 3.6 mg/kg bodyweight) and macaques with ketamine hydrochloride (Ketaset, 100mg/ml, Fort Dodge Animal Health Ltd., UK; 10mg/kg). SARS-CoV-2 Victoria/01/2020^26^ was prepared as described previously^24^. It was delivered to ferrets by intranasal instillation (1.0mL total, 0.5mL per nostril) diluted in PBS. A single dose of virus (5×10^6^ pfu/ferret) was delivered to Ad-GFP- (n=4) and FIV- (n=6) vaccinated ferrets. Macaques were challenged with 5 x 10^6^ delivered by the intratracheal route (2ml) and intranasal instillation (1ml total, 0.5ml per nostril). The schedule of euthanisation and sampling is shown in **Table 1**.

**Table 1.**
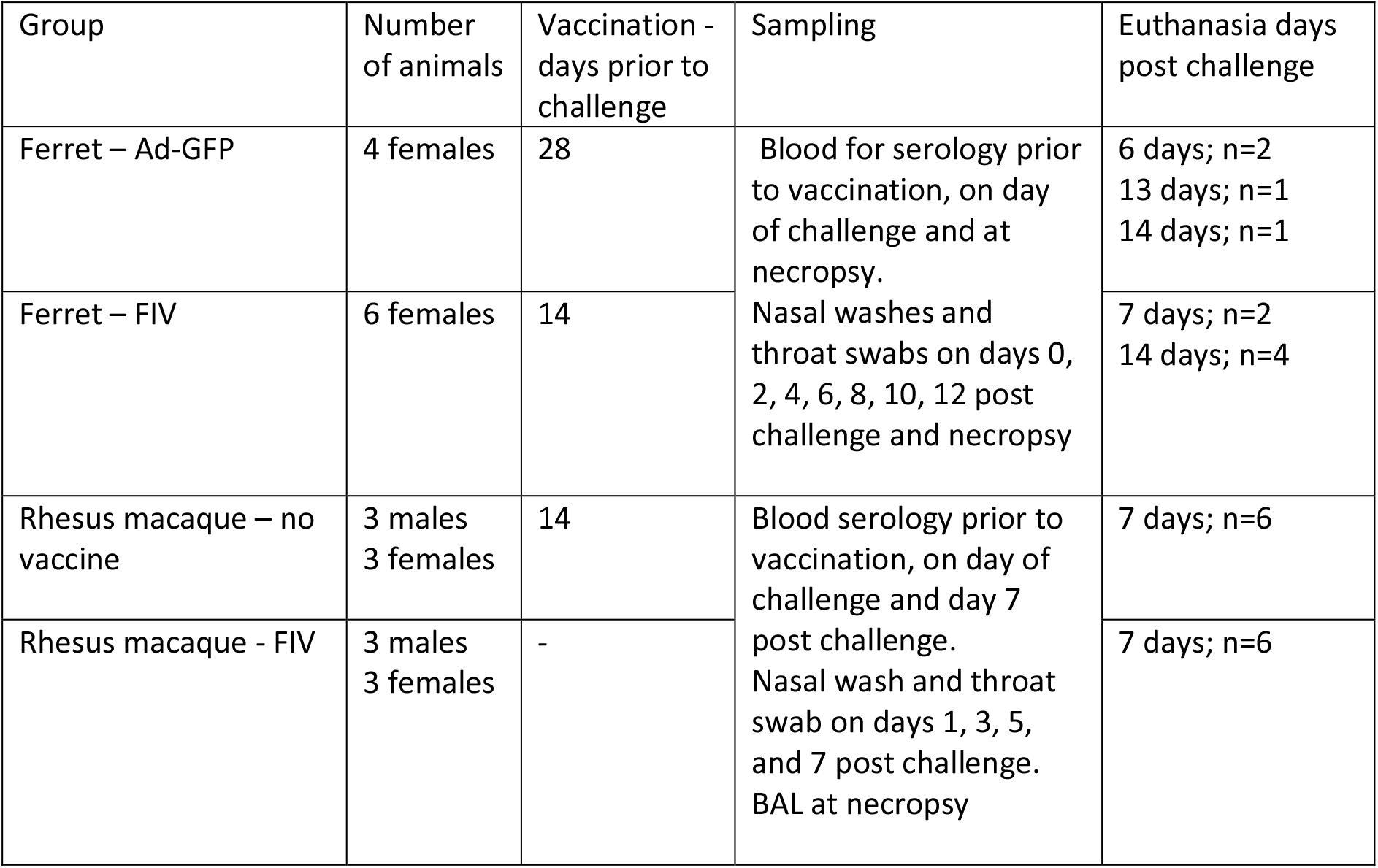
Experimental animal groups.

Nasal washes were obtained by flushing the nasal cavity with 2mL PBS. Throat swabs were collected using a standard swab (Sigma Virocult®) gently stroked across the back of the pharynx in the tonsillar area. Throat swabs were processed, and aliquots stored in viral transport media (VTM) and AVL at ≤ −60°C until assay. Clinical signs of disease were monitored as described previously^24 25^. The necropsy procedures were also as described previously^24 25^.

### SARS-CoV-2 virology

RNA was isolated from nasal wash and throat swabs. Samples were inactivated in AVL (Qiagen) and ethanol. Downstream extraction was then performed using the BioSprint™96 One-For-All vet kit (Indical) and Kingfisher Flex platform as per manufacturer’s instructions.

Reverse transcription-quantitative polymerase chain reaction (RT-qPCR) targeting a region of the SARS-CoV-2 nucleocapsid (N) gene was used to determine viral loads and was performed using TaqPath™ 1-Step RT-qPCR Master Mix, CG (Applied Biosystems™), 2019-nCoV CDC RUO Kit (Integrated DNA Technologies) and QuantStudio™ 7 Flex Real-Time PCR System. Sequences of the N1 primers and probe were: 2019-nCoV_N1-forward, 5’ GACCCCAAAATCAGCGAAAT 3’; 2019-nCoV_N1-reverse, 5’ TCTGGTTACTGCCAGTTGAATCTG 3’; 2019-nCoV_N1-probe, 5’ FAM-ACCCCGCATTACGTTTGGTGGACC-BHQ1 3’. The cycling conditions were: 25°C for 2 min, 50°C for 15 min, 95°C for 2 min, followed by 45 cycles of 95°C for 3 s, 55°C for 30 s. The quantification standard was in vitro transcribed RNA of the SARS-CoV-2 N ORF (accession number NC_045512.2) with quantification between 1 x 10^1^ and 1 x 10^6^ copies/μL. Positive samples detected below the limit of quantification were assigned the value of 5 copies/μL, whilst undetected samples were assigned the value of 2.3 copies/μL, equivalent to the assay’s lower limit of detection.

### ELISA to quantify anti-S, RBD and N IgG

A full length trimeric and stabilised version of the SARS-CoV-2 Spike protein (amino acids 1-1280, GenBank: MN MN908947) was developed by Florian Krammer’s lab as described^28^. Recombinant SARS-CoV-2 Receptor-Binding-Domain (RBD) (319-541) Myc-His was provided by MassBiologics. Recombinant SARS-CoV-2 Nucleocapsid phosphoprotein (GenBank: MN908947, isolate Wuhan-Hu-1) was expressed and purified from *Escherichia coli* as full-length nucleoprotein (amino acids 1-419) with a C-terminal 6xHis-Tag (Native Antigen Company). High-binding 96-well plates (Nunc Maxisorp, 442404) were coated with 50 μl per well of 2μg/mL Spike trimer, RBD or N in 1x PBS (Gibco) and incubated overnight at 4°C. The ELISA plates were washed five times with wash buffer (PBS 0.05% Tween 20) and blocked with 100 μL/well 5% FBS (Sigma, F9665) in PBS 0.1% Tween 20 for 1 h at room temperature. After washing, serum samples were serially diluted in 10% FBS in PBS 0.1% Tween 20 and 50 μl/well of each dilution was added to the antigen coated plate and incubated for 2 h at room temperature. Following washing, anti-ferret IgG-HRP (Novus Biologics, NB7224) diluted (1/1000) or anti-monkey IgG-HRP (Invitrogen PA-84631 diluted 1:10,000) in 10% FBS in 1X PBS/0.1% Tween 20 and 100 μl/well was added to each plate, then incubated for 1 h at room temperature. After washing, 1mg/mL O-phenylenediamine dihydrochloride solution (Sigma P9187) was prepared and 100 μL per well were added. The development was stopped with 50μL per well 1M Hydrochloric acid (Fisher Chemical, J/4320/15) and the absorbance at 490 nm was measured. OD1.0 titres were calculated using Softmax Pro 7.0.

### SARS-CoV-2 neutralisation assays

The plaque reduction neutralisation test (PRNT) was performed as described previously with ferret serum^24^. The microneutralisation assay was performed with macaque serum as described for human sera^29^.

### Isolation of Immune Cells

Similar to as described previously^24^ heparinised blood and spleens were removed for the isolation of immune cells; peripheral blood mononuclear cells (PBMCs) and splenocytes. The spleens were dissected into small pieces. Dissected spleen was dissociated using a gentleMACS. The tissue solution was passed through two cell sieves (100μm then 70μm) and then layered with Ficoll®-Paque Premium (GE Healthcare, Hatfield, United Kingdom). Density gradient centrifugation was carried out at 400g for 30 min on dissociated tissue and on whole blood. Buffy coats containing lymphocytes were collected and washed with medium by pelleting cells via centrifugation at 400 g for 10 min. The cells were counted using a vial-1 cassette and a Nucleocounter-200 before cryopreservation in 95% FCS/5% v/v DMSO. Cryopreserved cells were then frozen at −80°C in controlled rate freezer containers overnight, before transfer to liquid nitrogen (vapour phase).

### Interferon-gamma (IFN-γ) ELISpot Assay

An IFN-γ ELISpot assay was performed as described previously for ferrets^24^ and macaques^25^.

### Immunophenotyping

Whole blood immunophenotyping assays were performed using 50 μl of heparinised blood incubated for 30 min at room temperature with optimal dilutions of the following antibodies: anti-CD3-AF700, anti-CD4-APC-H7, anti-CD8-PerCP-Cy5.5, anti-CD95-Pe-Cy7, anti-CD14-PE, anti-HLA-DR-BUV395, anti-CD25-FITC (all from BD Biosciences, Oxford, UK); anti-CD127-APC (eBioscience); anti-γδ-TCR-BV421, anti-CD16-BV786, anti-PD-1-BV711, anti-CD20-PE-Dazzle (all from BioLegend); and amine reactive fixable viability stain red (Life Technologies); all prepared in brilliant stain buffer (BD Biosciences). Red blood cell contamination was removed using a Cal-lyse reagent kit as per the manufacturer’s instructions (Thermofisher scientific). BD Compbeads (BD Biosciences) were labelled with the above fluorochromes for use as compensation controls. Following antibody labelling, cells and beads were fixed in a final concentration of 4% paraformaldehyde solution (Sigma Aldrich, Gillingham, UK) prior to flow cytometric acquisition.

Cells were analysed using a five laser LSRII Fortessa instrument (BD Biosciences) and data were analysed using FlowJo (version 10, Treestar, Ashland, US). Immediately prior to flow cytometric acquisition, 50 μl of Truecount bead solution (Beckman Coulter) was added to each sample. Leukocyte populations were identified using a forward scatter-height (FSC-H) versus side scatter-area (SSC-A) dot plot to identify the lymphocyte, monocyte and granulocyte populations, to which appropriate gating strategies were applied to exclude doublet events and non-viable cells. Lymphocyte sub populations including T-cells, NK-cells, NKT-cells and B-cells were delineated by the expression pattern of CD3, CD20, CD95, CD4, CD8, CD127, CD25, CD16 and the activation and inhibitory markers HLA-DR and PD-1. GraphPad Prism (version 8.0.1) was used to generate graphical representations of flow cytometry data.

### Histopathology

The following samples from each ferret were fixed in 10% neutral-buffered formalin, processed to paraffin wax and 4 μm thick sections cut and stained with haematoxylin and eosin (HE); respiratory tract (left cranial and caudal lung lobes; 3 sections from each lung lobe: proximal, medial and distal to the primary lobar bronchus), trachea (upper and lower), larynx, tonsil, liver, kidney, spleen, mediastinal lymph node, and small (duodenum) and large intestine (colon). Nasal cavity samples were also taken and decalcified in an EDTA solution for 3 weeks before embedding. These tissues above were examined by light microscopy and evaluated subjectively. Three qualified veterinary pathologists examined the tissues independently and were blinded to treatment and group details and the slides randomised prior to examination in order to prevent bias. A semiquantitative scoring system was developed to compare the severity of the lung lesions for each individual animal and among groups. This scoring system was applied independently to the cranial and caudal lung lobe tissue sections using the following parameters: a) bronchial inflammation with presence of exudates and/or inflammatory cell infiltration; b) bronchiolar inflammation with presence of exudates and/or inflammatory cell infiltration; c) perivascular inflammatory infiltrates (cuffing); and d) infiltration of alveolar walls and spaces by inflammatory cells, mainly mononuclear. The severity of the histopathological lesions was scored as: 0=none (within normal limits), 1=minimal, 2=mild, 3=moderate, and 4=severe.

Tissue sections of both lung lobes, nasal cavity and gastrointestinal tract from animals culled at the early timepoint (day 6/7 post-challenge) were stained using the RNAscope *in situ* hybridisation (ISH) technique to identify SARS-CoV-2 RNA. Briefly, tissues were pre-treated with hydrogen peroxide for 10 min (room temperature), target retrieval for 15 min (98-101°C) and protease plus for 30 mins (40°C) (Advanced Cell Diagnostics). A V-nCoV2019-S probe (Cat No. 848561, Advanced Cell Diagnostics) was incubated on the tissues for 2 h at 40°C. Amplification of the signal was carried out following the RNAscope protocol using the RNAscope 2.5 HD Detection kit – Red (Advanced Cell Diagnostics).

In addition, immunohistochemistry was used to identify T cells (CD3^+^) in lung tissue sections. Samples were cut at 4μm onto adhesive slides and stained using the Leica Bond RxM (Leica Biosystems, Germany). Briefly, slides were dewaxed and treated with peroxide block for 5 min. Epitope retrieval was performed using Epitope Retrieval solution 2 (Leica Biosystems, Germany) for 20 min. A polyclonal rabbit anti-human CD3 antibody (1:200; Agilent Technologies Inc, CA) was applied for 15 min and used with Leica Polymer Refine Detection kit to complete the staining.

The following samples from each rhesus macaque were fixed, processed, cut and stained as described above: left cranial and caudal lung lobes, trachea, larynx, mediastinal lymph node, tonsil, spleen, liver, kidney, duodenum and colon.

For the lung, three sections from each left lung lobe were sampled from different locations: proximal, medial and distal to the primary lobar bronchus. A scoring system was used to evaluate objectively the histopathological lesions observed in the lung tissue sections^25^. The scores for each histopathological parameter were calculated as the average of the scores observed in the six lung tissue sections evaluated per animal.

Sections from the lung lobes, duodenum and colon were stained with RNAScope ISH as described above. For the lung sections, digital image analysis was carried out with Nikon NIS-Ar software in order to calculate the total area of the lung section positive for viral RNA.

### In-life imaging of macaques by computed tomography (CT)

CT scans were collected four weeks before vaccination and five days after challenge. CT imaging was performed on sedated animals using a 16 slice Lightspeed CT scanner (General Electric Healthcare, Milwaukee, WI, USA) in both the prone and supine position to assist the differentiation of pulmonary changes at the lung bases caused by gravity dependant atelectasis, from ground glass opacity caused by SARS-CoV-2. All axial scans were performed at 120 KVp, with Auto mA (ranging between 10 and 120) and were acquired using a small scan field of view. Rotation speed was 0.8 s. Images were displayed as an 11 cm field of view. To facilitate full examination of the cardiac and pulmonary vasculature, lymph nodes and extrapulmonary tissues, Niopam 300 (Bracco, Milan, Italy), a non-ionic, iodinated contrast medium, was administered intravenously (IV) at 2 ml/kg body weight and scans were collected immediately after injection and ninety seconds from the mid-point of injection.

Scans were evaluated by a medical radiologist expert in respiratory diseases, including in non-human primates^30^, blinded to the animal’s clinical status, for the presence of: disease features characteristic of COVID-19 in humans (ground glass opacity (GGO), consolidation, crazy paving, nodules, peri-lobular consolidation; distribution: upper, middle, lower, central 2/3, bronchocentric); pulmonary embolus and the extent of any abnormalities estimated (<25%, 25-50%, 51-75%, 76-100%).

### CT Score system

To provide the power to discriminate differences between individual NHP’s with low disease volume (i.e. <25% lung involvement), a score system was applied in which scores were attributed for possession of abnormal features characteristic of COVID in human patients (COVID pattern score) and for the distribution of features through the lung (Zone score). The COVID pattern score was calculated as sum of scores assigned for the number of nodules identified, and the possession and extent of GGO and consolidation according to the following system: Nodule(s): Score 1 for 1, 2 for 2 or 3, 3 for 4 or more; GGO: each affected area was attributed with a score according to the following: Score 1 if area measured < 1 cm, 2 if 1 to 2 cm, 3 if 2 −3 cm, 4 if > 3 cm and scores for each area of GGO were summed to provide a total GGO score; Consolidation: each affected area was attributed with a score according to the following: 1 if area measured < 1 cm, 2 if 1 to 2 cm, 3 if 2 −3 cm, 4 if > 3 cm. Scores for each area of consolidation are summed to provide a total consolidation score. To account for estimated additional disease impact on the host of consolidation compared to GGO, the score system was weighted by doubling the score assigned for consolidation. To determine the zone score, the lung was divided into 12 zones and each side of the lung divided (from top to bottom) into three zones: the upper zone (above the carina), the middle zone (from the carina to the inferior pulmonary vein), and the lower zone (below the inferior pulmonary vein). Each zone was further divided into two areas: the anterior area (the area before the vertical line of the midpoint of the diaphragm in the sagittal position) and the posterior area (the area after the vertical line of the mid-point of the diaphragm in the sagittal position). This results in 12 zones in total where a score of one is attributed to each zone containing structural changes. The COVID pattern score and the zone are summed to provide the Total CT score.

### ELISA to characterise ligand binding to formaldehyde-treated and untreated S trimer and RBD

Antigens and ligands. The S trimer expression plasmid was obtained from R Shattock and P. McCay (Imperial College London, UK), and expressed the Wuhan-Hu-1 sequence (NCBI Reference NC_045512.2) in the context of a functional S1S2 cleavage site, and with the addition of a C-terminal trimerization domain and 6x his and Myc tags. The RBD-Fc expression plasmid was also based on the Wuhan-Hu-1 sequence and was obtained from the Krammer lab (Mount Sinai, NY, USA). Proteins were expressed in 293F cells and purified by nickel column (Thermofisher) for S trimer and protein A column (Thermofisher) for RBD-Fc. Soluble ACE2-Fc was based on a published sequence^31^ and was obtained from H. Waldmann and manufactured by Absolute Antibody Inc, and RBD-binding mAbs CR3022^32^ and EY6A3^33^ were obtained from T. Tan and K-Y. Huang respectively, and were biotinylated using NHS-LC-biotin following manufacturer’s instructions (ThermoFisher).

For S trimer capture ELISA, high protein binding ELISA plates (PerkinElmer) were coated overnight at 4°C with anti-myc antibody 9E10 at 4 μg/mL. After washing and blocking in PBS/2% BSA/0.05% tween 20, S trimer at 1 μg/mL was added in 50μL/well in PBS/1% BSA/0.05% tween 20 (ELISA buffer, EB) for 2 h at room temperature. For RBD capture ELISA, high protein binding ELISA plates (PerkinElmer) were coated overnight at 4°C with rabbit anti-human IgG (Jackson Laboratories) at 5 μg/mL in PBS. After washing and blocking, RBD-Fc at 1 μg/mL was added in 50μL/well in EB for 2h at room temperature. After washing, wells requiring formaldehyde (FA) treatment were incubated with 50μL 0.02% methanol-free formaldehyde (ThermoFisher Scientific) in PBS for 72 h, and untreated wells with PBS for the same amount of time. After washing, S-captured plates were incubated with soluble ACE2-Fc and mAbs CR3022^32^ and EY6A^33^ in EB, and binding detected using donkey anti-human HRP (Jackson) at 1:5000 in 50μL EB. RBD-captured plates were incubated with biotinylated ligands soluble ACE2-Fc, CR3022 and EY6A, and binding detected using streptavidin-HRP (GE Healthcare) diluted 1:4000 in 50μL EB. Plates were incubated for 1 h at room temperature, washed, developed in 50μL TMB ELISA substrate (ThermoFisher) and the reaction stopped with 50μL 0.5M H2SO4. Absorbance at 450 and 570 nm was read on a SpectraMax M5 plate reader (Molecular Devices) and data analysed in GraphPad Prism v7.

### Molecular modelling

The structure shown is PDB 6lzg, and the views shown were generated in Pymol 2.3.5 (Schrodinger LLC).

## Results

### Characterisation of formaldehyde-inactivated vaccine (FIV)

Transmission electron microscopy (**Fig. 1**) showed that the washing and formaldehyde inactivation procedure resulted in virus particles that appeared similar to typical coronavirus morphology with a complete ring of peplomers/spikes on each particle (**Fig. 1A and B**). SDS-PAGE analysis (Supplementary Figure 1A) shows that the majority of protein bands seen in the medium only (lane 2) are also seen in the live wild-type virus (lane3) and the FIV (lane 4), indicating that these proteins are likely to be components of the culture medium, including FBS and host cell proteins. The medium-only protein species are visibly reduced in intensity following washing using the centrifugal concentrator in the FIV even though this was six-fold concentrated. Notably, protein species in the 62 to 100kDa range were almost absent in FIV. The Western blot analysis (**Supplementary Figure 1B and 1C**) confirms that both wild-type live virus and FIV react with antibodies to both SARS-CoV-2 spike-RBD and nucleocapsid. Western blot with NIBSC, SARS-CoV-2 neutralising, MERS convalescent serum (S3) standard has limited reactivity with proteins found in the medium (Supplementary Figure 1B). However, a virus-specific band corresponding to the spike-RBD band on the specific antisera blot is detected at approximately 160kDa in unwashed virus preparation and at 180kDa in FIV with the increase molecular weight likely to be a consequence of formaldehyde-fixation cross linking this protein.

**Fig. 1.**
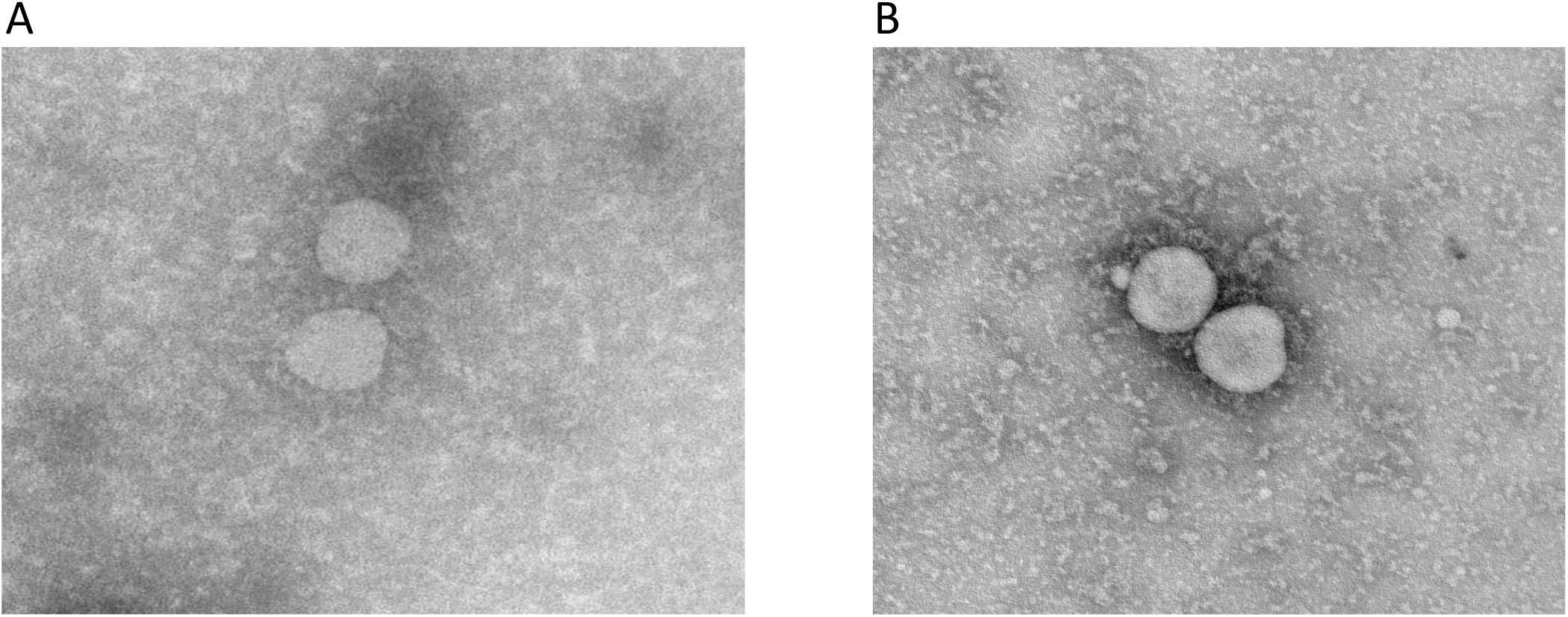
Representative transmission electron microscopy images of (A) the initial SARS-CoV-2 virus preparation and (B) following formaldehyde inactivation and washing to remove medium constituents.

### SARS-CoV-2 infection in control and FIV-vaccinated ferrets

Ferrets vaccinated with either a recombinant adenovirus expressing GFP (Ad-GFP) or FIV were challenged intranasally with 1mL of Victoria/1/2020 SARS-CoV-2 at 5 x 10^6^ PFU^26^. The sampling schedule is shown in **Table 1**. The high titre stock of challenge virus was prepared (passage 3), and quality control sequencing showed it was identical to the original stock received from the Doherty Institute and did not contain a commonly reported 8 amino acid deletion in the furin cleavage site^34^. Both groups displayed similar viral genome copies in nasal wash samples which continued to be detected until the end of the experiment at day 15 pc (**Fig. 2A**). Similar to previous studies^24^ the peak in viral RNA shedding was seen between day 2 and 4 pc for all ferrets in both groups. The majority of virus detected in nasal wash occurred between challenge and day 8 pc. Interestingly, viral RNA detected in nasal wash and throat swab samples was also shown to be approximately 3-fold higher in the FIV group at day 2 pc. A similar trend was seen in both groups during the first week after challenge although the RNA genome copies measured were substantially lower in the throat swab than in the nasal wash samples (**Fig. 2B**). Overall, there were no significant differences between groups in virus shedding from either the nose or throat. Neither group of animals showed weight loss due to the infection (**Supplementary Fig. 2**). The apparent difference in weight gain in the FIV-vaccinated group was due to the necropsy-sampling of the two lightest animals from this group on day 7. No fever was seen at any time in either group of ferrets post infection (**Supplementary Fig. 2**).

**Fig. 2.**
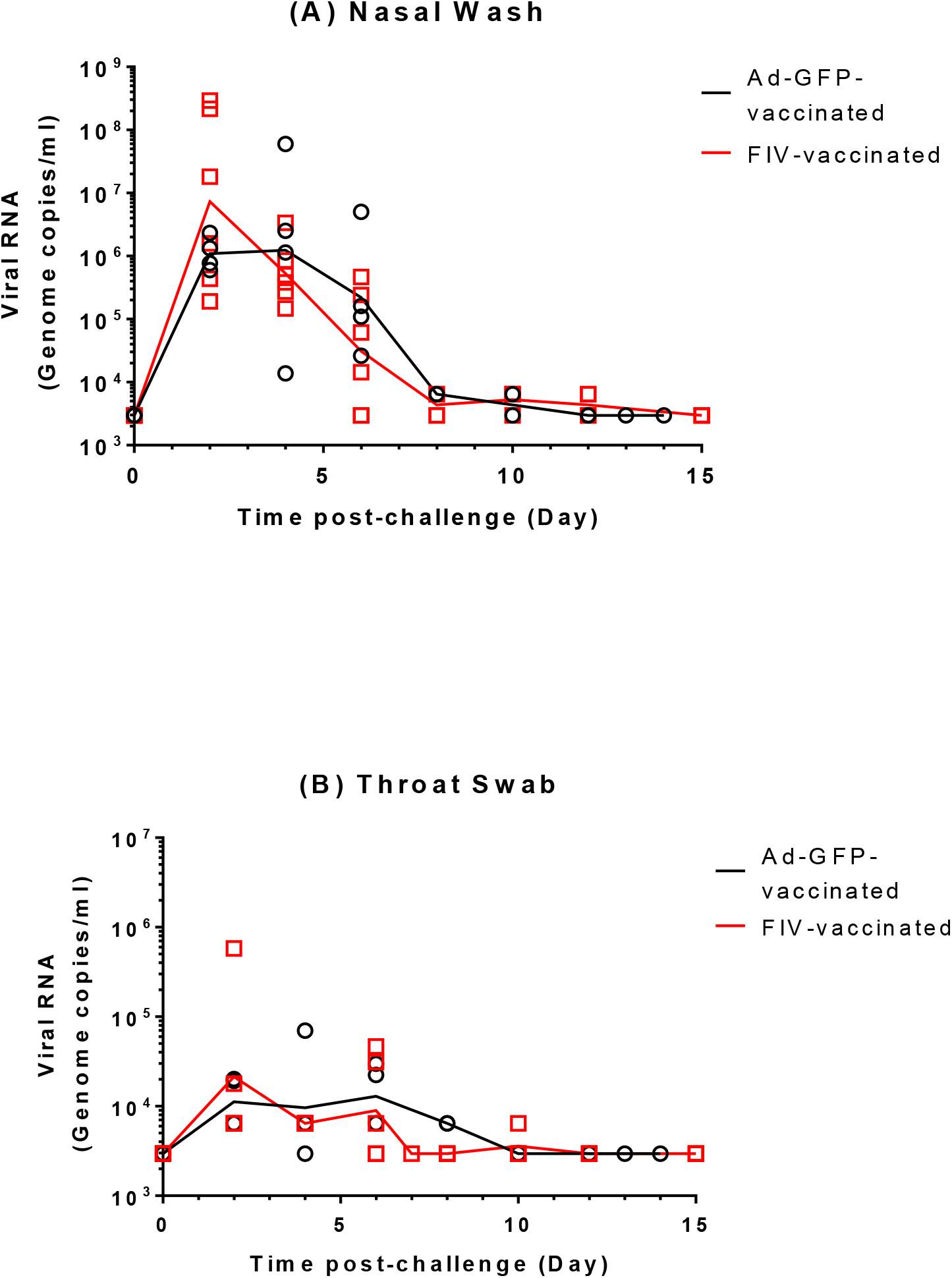
Detection of SARS-CoV-2 RNA in ferret respiratory samples. Viral RNA in Ad-GFP and FIV-vaccinated ferrets was quantified by RT-PCR in (A) nasal washes and (B) throat swabs. Lines plotted are the geometric mean genome copies per mL.

### Pathology following SARS-CoV-2 infection in ferrets

We performed sequential culls on days 6-7 and 13-14 in order to study the potential for VED during and after resolution of SARS-CoV-2 infection. The lung histopathology scores for individual Ad-GFP- and FIV-vaccinated ferrets are shown in the heat map in **Fig 3A**. Samples were obtained from 2 animals from each group early in the infection (Ad-GFP at day 6 pc and FIV at day 7 pc). The remaining animals were euthanised at days 13-14 pc (Ad-GFP) and day 15 pc (FIV). All assessments, including bronchiolar, bronchial and interstitial infiltrates together with perivascular cuffing, were scored as minimal or mild in the Ad-GFP-vaccinated animals with a greater number of mild or moderate scores in the FIV-vaccinated ferrets at the early time point (6/7 days pc). One animal from the Ad-GFP group showed mild lesions compatible with acute bronchiolitis and perivascular/peribronchiolar cuffing (**Fig. 3F,G**). The other animal from this group showed only occasional minimal bronchiolar infiltrates. Both animals from the FIV group at 7 days pc showed more remarkable changes, with mild to moderate bronchiolitis (infiltrates within the bronchioles and occasionally bronchi) and inflammatory foci within the parenchyma (**Fig. 3B**). Moreover, perivascular cuffing was observed frequently (**Fig. 3C**), with the infiltrates being mostly mononuclear cells, including CD3^+^ T lymphocytes identified by immunohistochemistry (IHC) staining (**Fig. 3D**). Occasionally, neutrophils and eosinophils were also present (**Fig. 3C, insert**). The cuffing also affected numerous airways (**Fig. 3C**). Due to the small numbers of animals, the differences in scores observed between FIV and Ad-GFP-vaccinated groups did not reach significance.

**Fig. 3.**
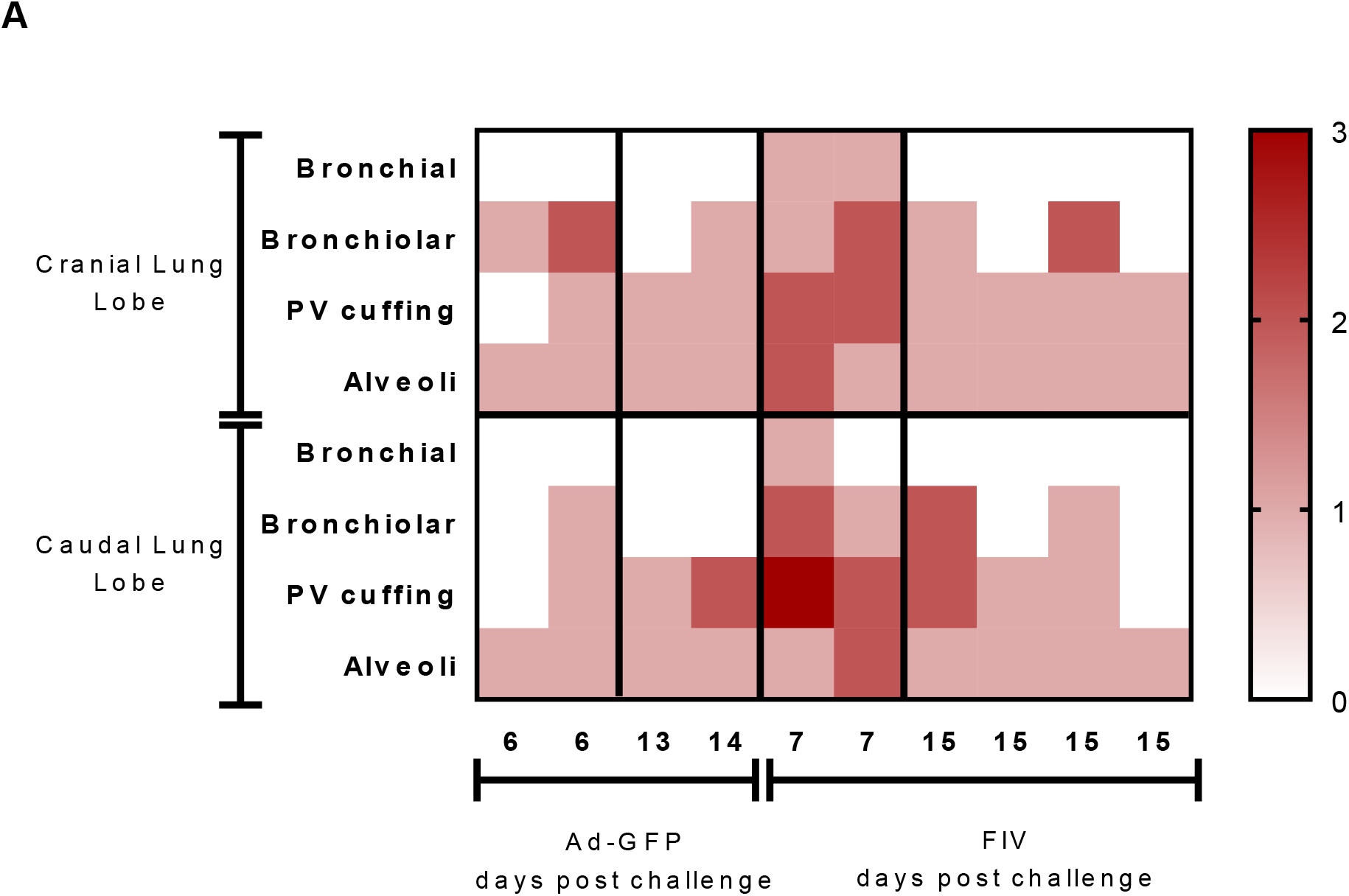
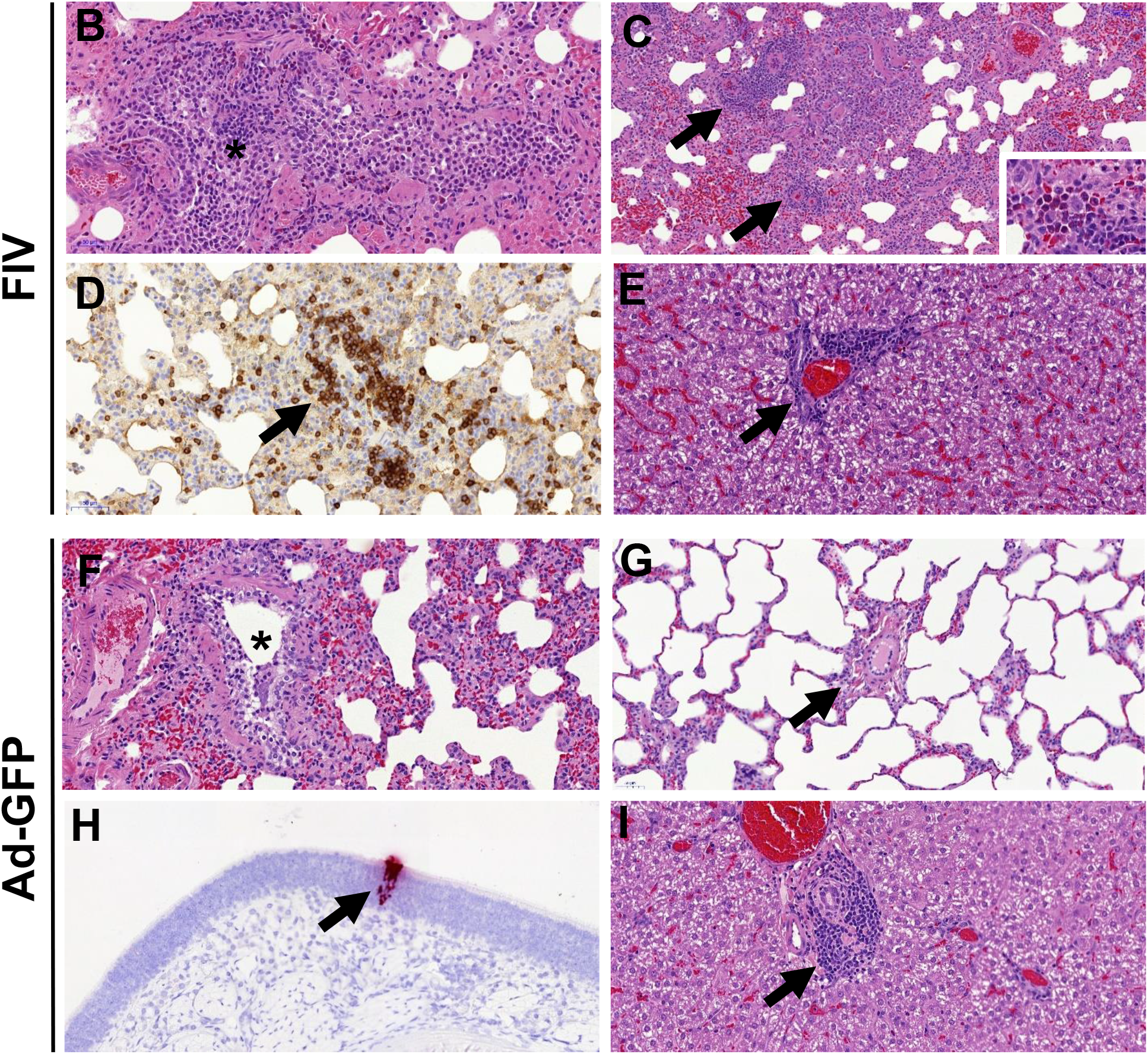
A: Heatmap showing the individual lung histopathology scores for each ferret and parameter following FIV- or Ad-GFP-vaccination, and challenged with SARS-CoV-2 and culled at 6/7 days and 13/15 days pc. Histopathology of FIV- (B-E) and Ad-GFP (F-I)-vaccinated ferrets. **B.** Inflammatory infiltrates within a bronchiole (*), with abundant mononuclear cells but also some neutrophils and eosinophils. H&E, 200x. **C.** Multiple inflammatory infiltrates surrounding blood vessels (perivascular cuffing, arrows). H&E, 100x. The infiltrates are composed mostly of macrophages and lymphocytes, but abundant eosinophils can also be observed in some areas within the infiltrates (insert; H&E, 400x). **D.** A perivascular cuff (arrow) with abundant mononuclear cells, many of them identified as CD3^+^ T lymphocytes. IHC, 200x. **E.** Periportal mononuclear inflammatory infiltrate in the liver (mild multifocal hepatitis). H&E, 400x. **F.** Mild inflammatory infiltrate within a bronchiole (*). H&E, 200x. **G.** Blood vessel (arrow) within the lung parenchyma not showing any perivascular cuffing. H&E, 200x. **H.** ISH detection of SARS-CoV-2 RNA in a small focus of epithelial and sustentacular cells within the nasal cavity. ISH, 200x. **I.** Periportal mononuclear inflammatory infiltrate in the liver (mild multifocal hepatitis). H&E, 400x.

In contrast, at 13-15 days pc, the lesions observed were minimal to mild with no obvious differences between groups (**Fig. 3A**).

RNAScope ISH technique was used to detect viral RNA in lung and nasal cavity tissue sections. Only very few occasional scattered cells were found positive to viral RNA in the lung at day 6/7, which were within the alveolar walls and not related to the presence of lesions. No differences were observed between groups. Viral RNA was also found only as small foci of positive cells (epithelial and or sustentacular) within the olfactory and respiratory mucosa in only one animal from the Ad-GFP group at day 6 pc (**Fig. 3H**).

No obvious lesions were observed in any other organ except for the liver, which showed a variable degree of multifocal hepatitis, mild to moderate in all animals (**Fig 3E, I**).

### Immune responses to FIV in ferrets

Ad-GFP-vaccinated animals showed no immune responses before SARS-CoV-2 challenge (**Fig 4**). FIV-vaccinated animals produced significant increases in IgG after vaccination against SARS-CoV-2 spike and spike receptor binding domain (RBD). The response to nucleoprotein (N) was not significant. Modest rises (GMT=89, p=0.002) in neutralising antibody titres were seen in sera from the FIV-vaccinated animals with a rapid rise in neutralising antibody titres after challenge indicating immune priming by the FIV. At termination, the GM titres for FIV-vaccinated animals were 5356 and 453 for the two remaining Ad-GFP-vaccinated animals. ELISpot assays applied to splenocytes isolated on day 6 pc (Ad-GFP-vaccinated) or day 7 pc (FIV-vaccinated) show that animals vaccinated with FIV made more pronounced responses to whole live virus, membrane, and nucleocapsid peptide pools with both groups showing similar low responses to virus spike peptides (**Supplementary Fig. 3A**). This early interferon gamma response post-infection is consistent with an anamnestic response in the FIV-vaccinated animals following priming with viral antigens. The lack of cellular immune response to spike is interesting and might indicate that FIV promotes a skewed immune response. Responses measured in PBMCs were much lower than those seen in splenocytes.

**Fig. 4.**
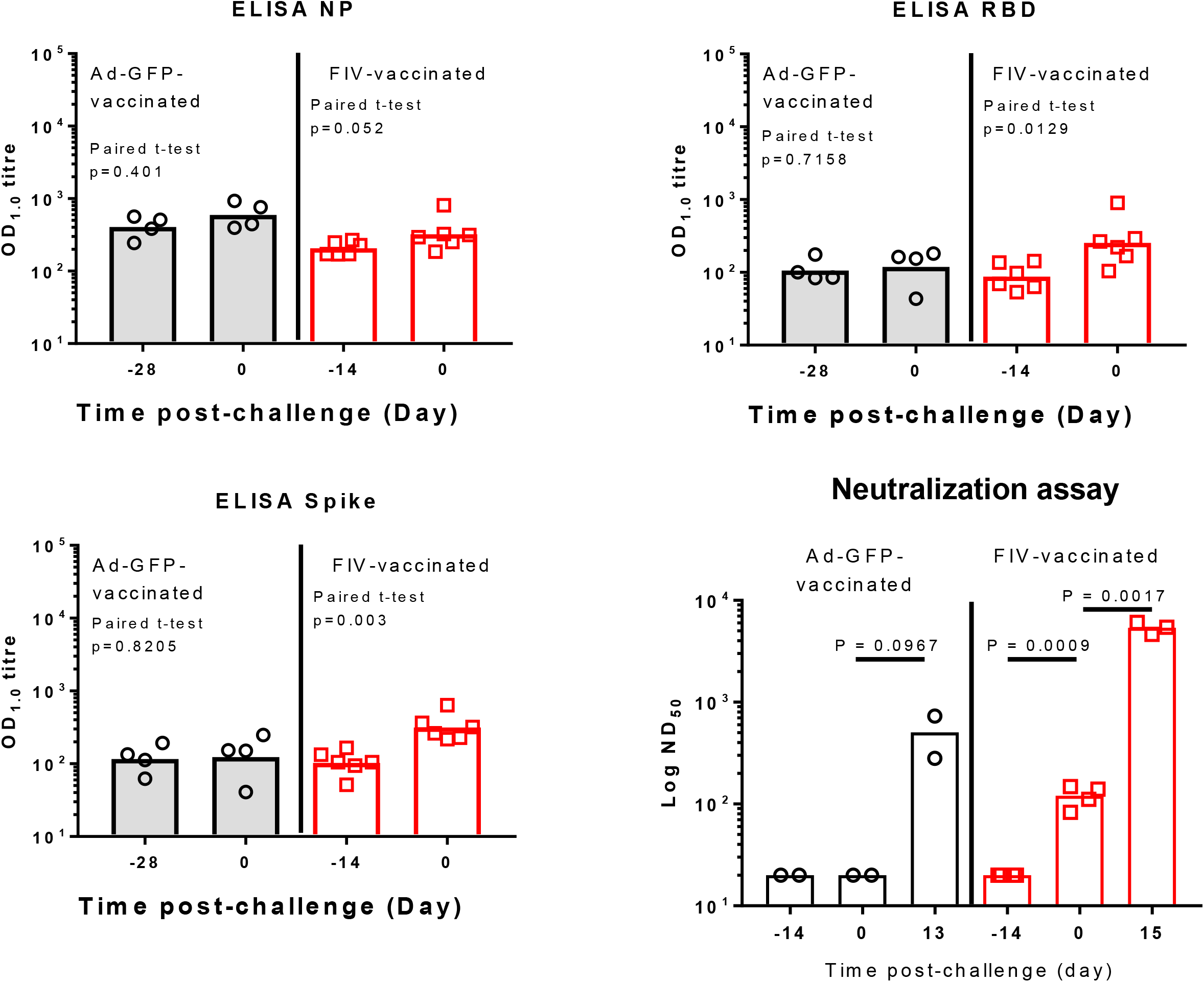
Serological response to Ad-GFP and FIV in ferrets. IgG was quantified by ELISA to recombinant nucleocapsid protein (NP), receptor binding domain (RBD) and full-length trimeric and stabilised spike protein (Spike). Bars are geometric mean titre. The significance of any difference from pre-to post-vaccination is shown, determined by a paired t-test. The plaque reduction neutralisation 50% titre (PRNT_50_) is also shown with samples obtained pre- and post-vaccination and following SARS-CoV-2 challenge. Bars are geometric mean PRNT_50_ titre.

### SARS-CoV-2 infection in control and FIV-vaccinated rhesus macaques

Following infection with 5 x 10^6^ PFU Victoria/1/2020 SARS-CoV-2 given in 2ml by the intratracheal route and 1ml intranasally, viral RNA was quantified in nasal wash and throat swab samples. Viral RNA was detected in both samples during the experiment with peak values detected by day three and a decline thereafter **(Fig 5A and B)**. There was no difference between viral copies detected in macaques that received FIV or no vaccine. Bronchioalveolar lavage (BAL) was obtained on necropsy at day 7 and lower geometric mean viral RNA copies per ml (*p*=0.168) were measured in macaques that received FIV than no vaccine controls **(Fig 4C)**. No significant changes in body temperature (**Supplementary Fig. 2C**) where observed. Slight weight loss was observed in both groups (**Supplementary Fig. 2D**), but no adverse clinical signs were recorded for any macaque despite frequent monitoring during the study period.

**Fig. 5.**
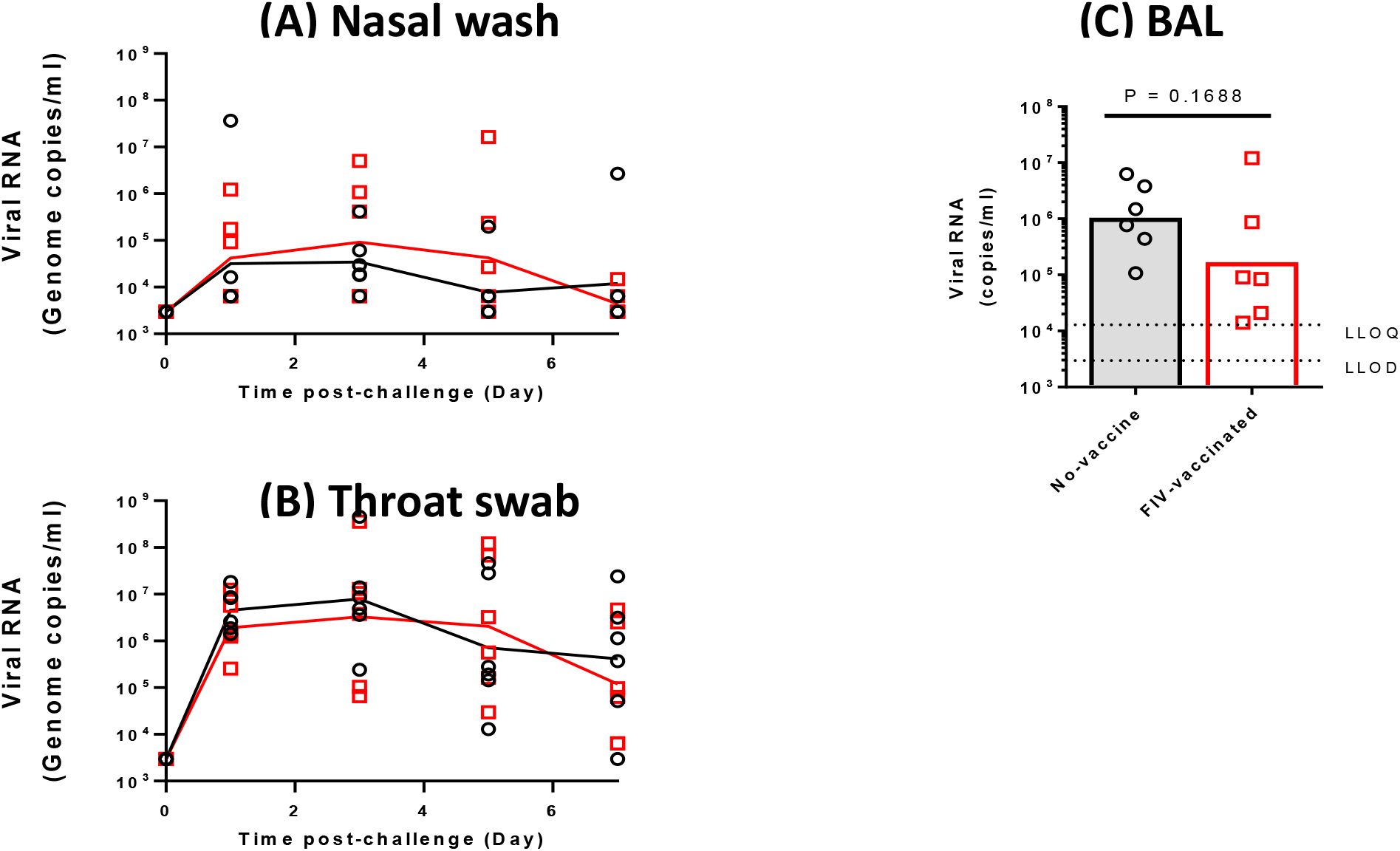
Detection of SARS-CoV-2 RNA in macaque respiratory samples. Viral RNA in unvaccinated and FIV-vaccinated macaques was quantified by RT-PCR in (A) nasal washes, (B) throat swabs and (C) bronchiolar lavage. Lines plotted are the geometric mean genome copies per mL.

Images from CT scans collected five days after challenge were examined by an expert thoracic radiologist with experience of non-human primate prior CT interpretation and human COVID-19 CT features, blinded to the clinical status. Pulmonary abnormalities that involved less than 25% of the lung and reflected those characteristics of SARS-CoV-2 infection in humans were identified in three of the FIV group and five of the unvaccinated group. Where reported, disease was predominantly bilateral (two of three FIV, five of six unvaccinated) with a similar peripheral distribution through the lung lobes reported in the FIV vaccinated and unvaccinated macaques. Ground glass opacity was observed in all the macaques showing abnormal lung structure, with the exception of one FIV-vaccinated animal in which consolidation was identified. Other features characteristic of human COVID (reverse halo, perilobular, nodules, pulmonary embolus) were not observed in any of the macaques in either group. Evaluation of pulmonary disease burden using a scoring system designed to discriminate differences between individual macaques with low disease volume revealed a non-significant trend (p = 0.1364) for a reduction in the total CT score in the FIV group compared to the scores attributed to macaques in the unvaccinated group (**Fig. 6D**). Similarly, the FIV vaccine reduced both the amount of abnormalities induced (pattern score) and distribution of disease (zone score) (**Supplementary Fig. 4 H and I**).

**Fig. 6.**
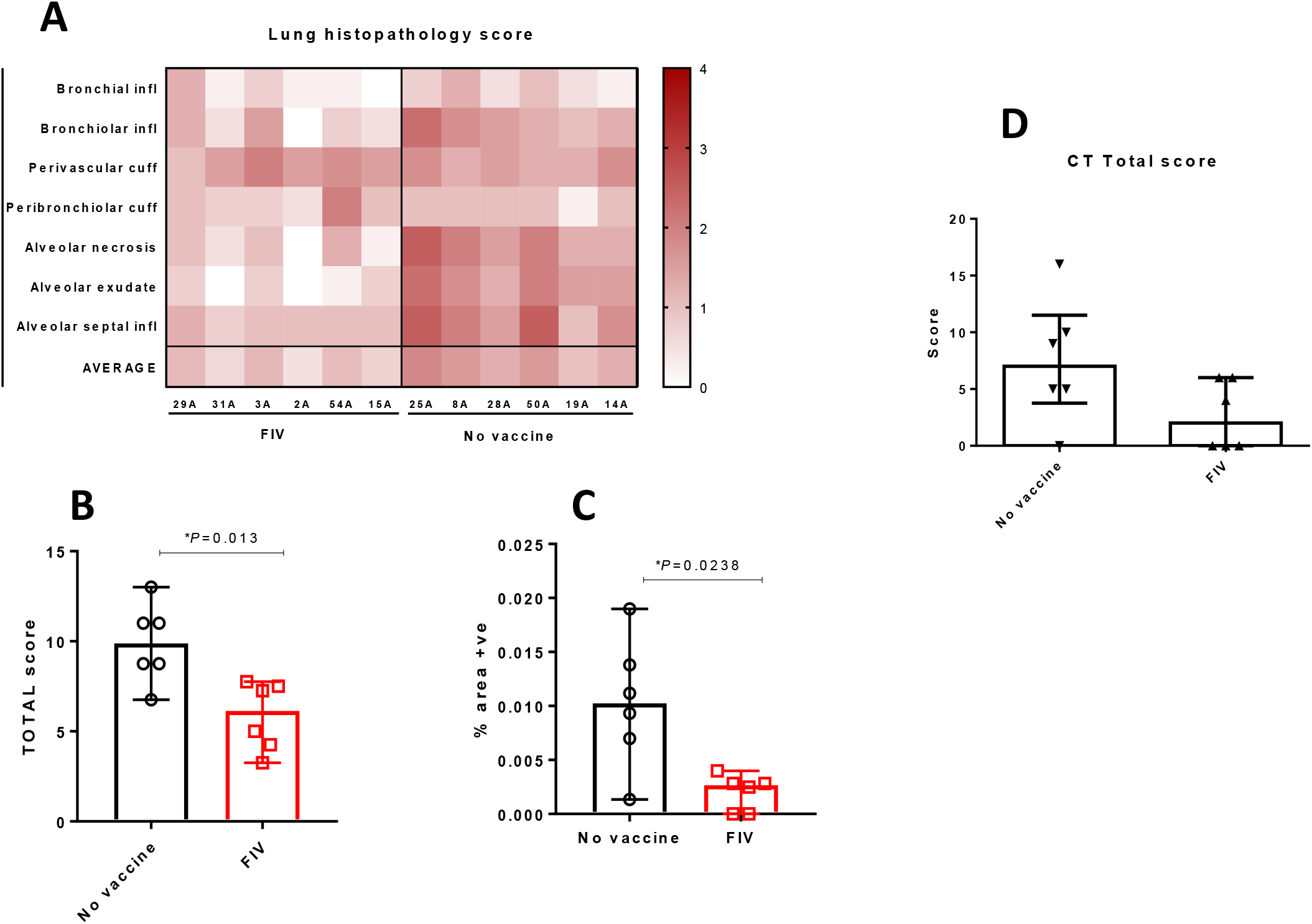
Histopathology and CT scan analysis in rhesus macaques. A) Heatmap showing the individual lung histopathology scores for each animal and parameter, and the AVERAGE severity for each animal. B) Total histopathology scores in non vaccinates and FIV animals, showing a significant reduction in severity in the FIV group (*P*=0.013, Mann-Whitney U-tests; Boxes and whiskers show median +/- 95% C.I.). C) Percentage of area positively stained with ISH RNAScope (viral RNA) in non vaccinates and FIV animals, showing a significant reduction in the FIV group (*P*=0.0238, Mann-Whitney U-tests; Boxes and whiskers show median +/- 95% C.I.). D) Plot shows the total CT scores in non-vaccinated and FIV-vaccinated animals showing a non-significant trend for reduction in severity in the FIV group (p = 0.1364 Mann-Whitney U-test); Box plots show the experimental group median with +/- IQR indicated by box whiskers, symbols show scores measured in individual animals.

### Pathology following SARS-CoV-2 infection in rhesus macaques

Pathological changes were found in the lungs of all SARS-CoV-2-infected macaques and consisted of multiple areas of mild to moderate bronchiolo-alveolar necrosis, inflammatory cell infiltration and type II pneumocyte proliferation. Mild perivascular and peribronchiolar cuffing was also observed. The lung pathology scores for individual macaques are shown in **Fig. 6** and with milder pathological changes observed in the FIV-vaccinated macaques. The total pathology score for the no vaccine group was greater than the FIV group (**Fig. 6B**, *p*=0.013). RNAScope analysis of the percentage of area positively stained for SARS-CoV-2 RNA showed a greater lung area infected in the no vaccine group than FIV-vaccinated macaques (**Fig. 6C**, *p* = 0.0238).

### Immune responses to FIV in rhesus macaques

Serum from control macaques obtained on the day of challenge did not show any N, RBD or S-specific IgG but rises in RBD and S-specific IgG were detected in serum from the FIV-vaccinated macaques (Fig. 7). FIV-vaccinated animals also showed a rise in neutralising antibody titre on the day of challenge (*p*=0.0287). Both groups showed a rise in neutralising antibody titre 7 days following challenge (**Fig. 7**). Similarly, on the day of challenge, a higher frequency of spike-specific IFNγ-secreting cells was measured by ELISPOT assay in the FIV group compared to that determined in the unvaccinated group suggesting the induction of a modest but significant (*p* = 0.0433) SARS-CoV-2-specific cellular response (**Supplementary. Figure 3B**). The trend reversed six to eight days after challenge when frequencies were assessed at the end of the study, with higher frequencies of spike-specific IFNγ-secreting cells measured in the unvaccinated group compared to the FIV group in both PBMC and spleen cells.

**Fig. 7.**
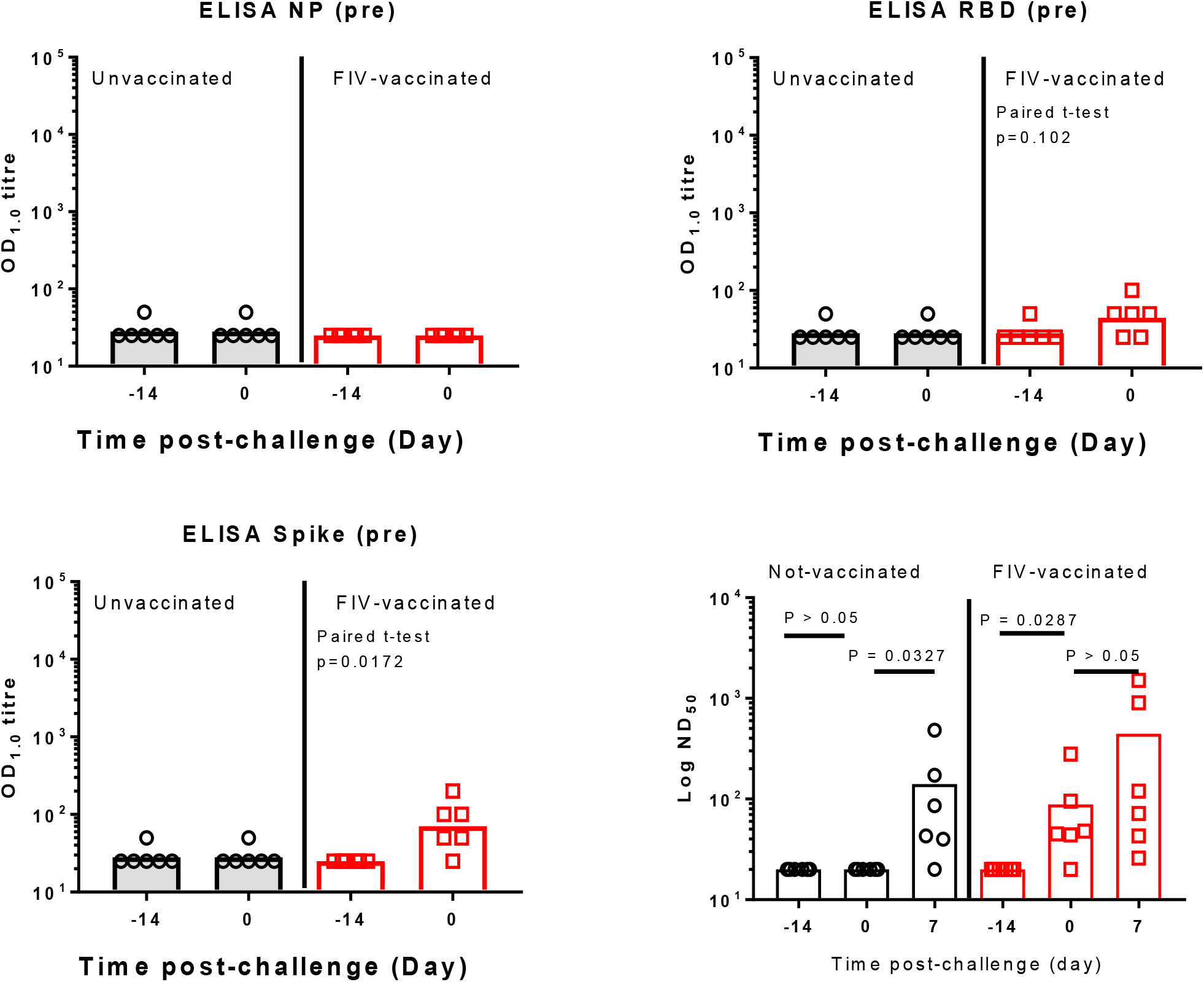
Serological response in unvaccinated and FIV-vaccinated macaques. IgG was quantified by ELISA to recombinant nucleocapsid protein (NP), receptor binding domain (RBD) and full-length trimeric and stabilised spike protein (Spike). Bars are geometric mean titre. The significance of any difference from pre-to post-vaccination is shown, determined by a paired t-test. The micronutralisation 50% titre (ND_50_) is also shown with samples obtained pre- and post-vaccination and following SARS-CoV-2 challenge. Bars are geometric mean ND_50_ titre.

Immunophenotyping flow cytometry assays were applied to whole blood samples collected immediately prior to and 14 days after FIV vaccination, as well as at days three and seven after SARS-CoV-2 challenge to explore potential vaccine-induced changes in the cellular immune compartment that might influence the course of the immune response following infection (**Sup figure 3**). CD4 and CD8 T-cells expressing the immune checkpoint signalling receptor PD-1 increased significantly following FIV vaccination (*p* = 0.03), with further significant increases observed in PD-1 expressing CD4 T-cell populations following SARS-CoV2 infection in both FIV vaccinated and unvaccinated groups (both *p* = 0.03) (SF3 D). Similarly, FIV vaccination led to a significant increase in CD4 regulatory T-cells expressing CD25 and CD127 (SP3 E), indicating that alongside the proinflammatory cellular response evident in the antigen-specific IFNγ ELISPOT profiles, FIV vaccination also induced T-cell populations with a more tolerogenic phenotype.

### Effects of formaldehyde on SARS-CoV-2 spike

Formaldehyde treatment of SARS-CoV-2 virus will cross link viral proteins, of which the S glycoprotein trimer is the target of most neutralising antibodies. Cross-linking of S may modify its antigenicity, potentially altering elicitation of neutralising antibodies. To analyse the effects of formaldehyde on the S trimer and the isolated receptor binding domain (RBD), soluble antigens were captured onto ELISA plates using either anti-Myc tag (S-Myc) or anti-Fc (RBD-Fc) respectively to maintain their native conformation, and treated or not with formaldehyde using the same protocol as for inactivation of whole virus: 0.02% for 72 h at room temperature. Samples were then tested for binding of RBD ligands, either soluble (s)ACE2-Fc or the RBD binding monoclonal antibodies (mAbs) CR3022 and EY6A which interact with RBD surfaces non-overlapping the ACE2 binding site. Binding curves revealed that ligands binding to formaldehyde-treated S protein gave substantially lower maximum binding than that to the untreated S counterpart (**Fig. 8A**). Area under the curve (AUC) analysis revealed that binding was significantly reduced for sACE2-Fc and CR3022, and had a trend to reduction for EY6A (**Fig. 8B**). Interestingly, the reduction for sACE2 and CR3022 was almost precisely 2-fold, suggesting either that the formaldehyde treatment had reduced the binding activity of a subset of RBD domains, or that half of the formaldehyde-treated S trimers were in a non-RBD available conformation. To differentiate between these two possibilities, we tested binding directly to the isolated untreated or formaldehyde-treated RBD (**Fig. 8C**). Strikingly, formaldehyde treatment had no effect on RBD-ligand binding, with AUC analysis showing near identical values for formaldehyde-treated and untreated RBD (**Fig. 8D**). It therefore seems most likely that formaldehyde treatment is stabilising a population of S trimers in the ‘RBD down’ conformation which would be unable to engage ACE2. Indeed, recent cryo-EM structures imply that the S trimer is 50% one RBD up and 50% all RBD down at equilibrium^35^. Therefore, cross-linking of a population of trimers would most likely fix this 1:1 equilibrium, allowing only half of the trimers to expose the one-up RBD. By contrast, untreated S trimer would be free to sample both conformations, allowing progressive ACE2 occupancy to maximum of the one RBD-up trimer conformation over time. This would explain the 2-fold decrease in occupancy of formaldehyde-treated S trimers shown above. To further interrogate this, we modelled the location of lysine and arginine residues, the side chains of which are targets for formaldehyde attack, in the S trimer and RBD structures. **Fig. 6E** focuses on the location of lysines and arginines in the RBD-down S trimer, revealing a large number proximal to the RBD-trimer interface that might be cross-linked to prevent RBD movement. **Fig. 6F** shows that whilst there are lysine and arginine residues proximal to the RBD-ACE2 binding interface (green), there are none within the interface, implying that formaldehyde treatment would not directly affect RBD-ACE2 binding. This modelling is therefore consistent with the idea that formaldehyde cross-linking will lock 50% of trimers into the all RBD-down state, reducing access of the RBD to ligand binding and B cell recognition. Such modified antigenicity would probably translate into reduced RBD immunogenicity, reducing antibody production against this neutralising antibody-eliciting surface.

**Fig. 8.**
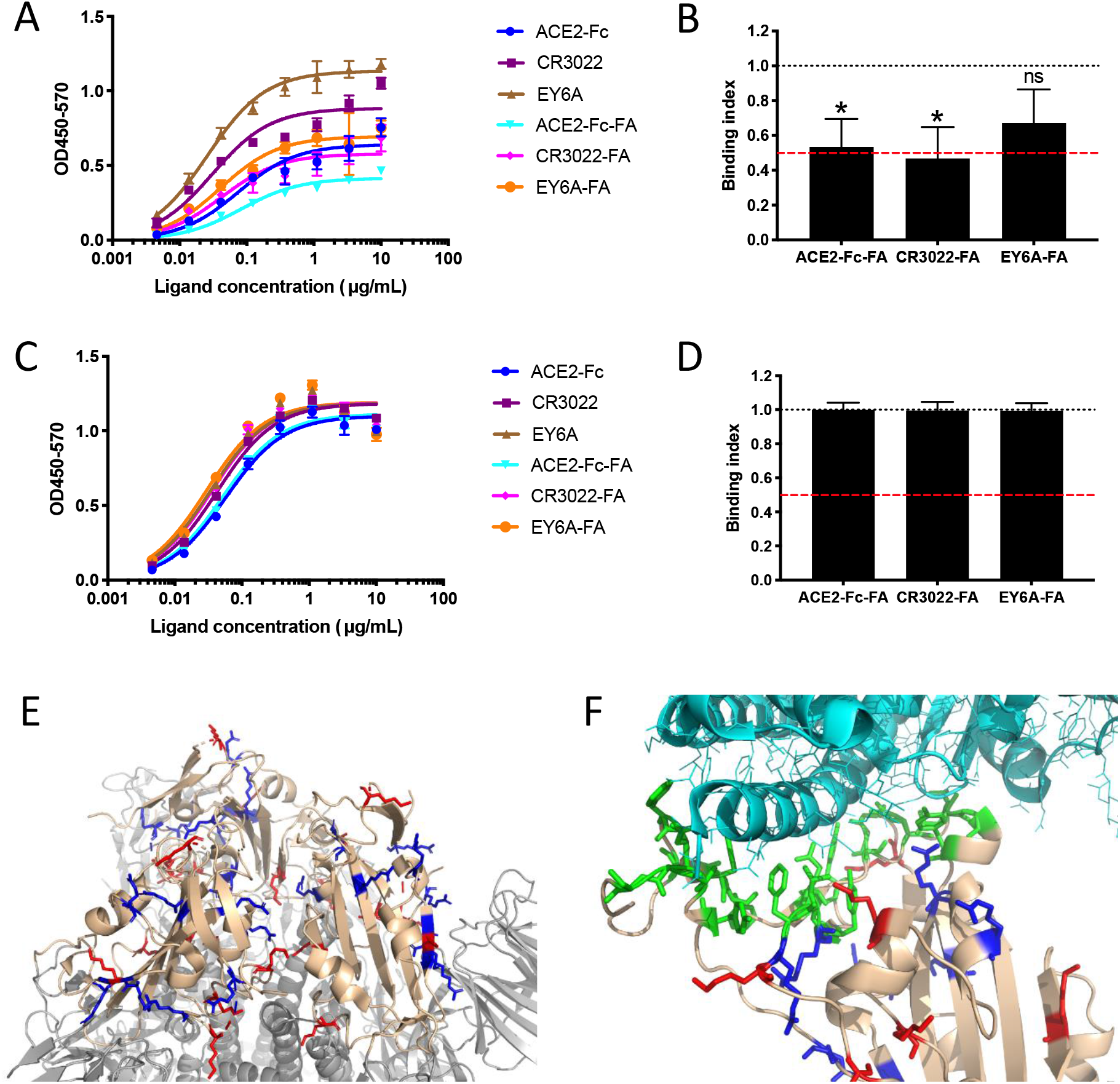
Formaldehyde treatment reduces S trimer binding to ACE2. A) Representative ELISA with S trimer captured onto the ELISA plate using anti-Myc mAb 9E10, untreated or treated with formaldehyde (FA) for 72 h at room temperature prior to addition of ligands at the concentrations shown and ligand detection using anti-human Fc-HRP. B) Area Under the Curve (AUC) analysis was used to determine the binding index, where 1=equivalent binding between untreated and FA-treated S trimer, and 0=zero binding to formaldehyde-treated trimer. The red line represents 50% binding. n=means of pooled data from 3 independent ELISAs, each performed in triplicate. *p<0.05 compared to unmodified condition, Student’s t-test after normality testing, ns=not significant. C) Representative ELISA with RBD-Fc captured onto ELISA plate with anti-human-Fc, untreated or treated with FA as for the S trimer. D) AUC analysis was used to determine the binding index as for S trimer above, n=means of pooled data from 3 independent ELISAs, each performed in triplicate. Student’s t-test after normality testing revealed no significant differences. E) Model prepared from PBD6LZG with 3 RBD ‘down’. RBD rendered in beige, S trimer in grey, lysines in red, arginines in blue. Close up of RBD-ACE2 interface with RBD in beige, ACE2 in light cyan, binding site in green, lysines in red, arginines in blue.

## Discussion

Rapid development of vaccines to prevent COVID-19 disease is in progress^2^. Safety is of primary importance for vaccines that are administered to healthy people. Thus vaccines must be thoroughly assessed for reactogenicity and longer term safety^36^, but also to ensure they do not cause any enhancement of disease^37^. Vaccine-enhanced disease (VED) which can be mediated by antibody-dependent enhancement (ADE) have been described for respiratory virus vaccines. The potential risks from COVID-19 vaccines have been described by Graham^38^. Enhanced disease can potentially be mediated by antibodies that bind virus without neutralising activity, which cause disease through increased viral replication, or via formation of immune complexes that deposit in tissues and activate complement pathways associated with inflammation^12^. T helper 2 cell (Th2)-biased immune responses have also been associated with ineffective vaccines that can lead to enhanced disease following infection^39 40^. In order to understand the risks of VED caused by SARS-CoV-2 vaccines it will be of great benefit to produce a positive control vaccine that enhances disease in an animal model following challenge so this endpoint can be defined and the mechanisms can be understood and avoided in vaccines prepared for human use.

As examples of VED have been observed following the use of a formaldehyde inactivated vaccine, e.g. RSV^41^ and measles^16^, we have prepared a killed SARS-CoV-2 vaccine by formaldehyde-fixation of the virus. Formaldehyde-fixed SARS-CoV-1 vaccines have been shown to induce enhanced disease^18 19 21 42^ although the mechanism for this is not understood. It has been suggested that non-viral components of formaldehyde-inactivated preparations, such as cellular components and debris or medium constituents may also play a role in enhanced disease. In a cotton rat model of RSV VED, cell culture contaminants were a major driver of lung pathology which was exacerbated by the formaldehyde-inactivated vaccine and RSV challenge^43^. The SARS-CoV-2 FIV prepared in this study also contained cell and medium components although the content was significantly reduced by washing using a centrifugal concentrator. Another factor in the FIV vaccine design was to use Alhydrogel as an adjuvant which is known to induce Th2-biased immune responses^44^. In addition, as a sub-optimal immune response has been suggested to be associated with VED^14^, we chose to challenge the ferrets and rhesus macaques 14 days after intramuscular delivery of a single dose of FIV.

The SARS-CoV-2 infection in the Ad-GFP- or FIV-vaccinated ferrets followed a similar course to that observed in our previous study^24^ with peak viral RNA shedding between 2 and 4 days pc. It was interesting to note that higher viral loads were detected in the upper respiratory tract of FIV-vaccinated animals at day 2 pc but after this sampling time, very similar genome copy values were obtained. There were no differences in temperature, weight (Supplementary figure 2) or any other clinical signs between the two groups.

Consistent with the higher viral load at day 2, the lung histopathology from the two FIV-vaccinated animals necropsied at day 7 was more severe than the two Ad-GFP-vaccinated animals necropsied at day 6. The semiquantitative scoring system was used to discriminate the severity of lesions between animals and groups. Even though the number of animals was small, and although lung pathology was not severe in any case, we observed some unique differences in the FIV-vaccinated ferrets. At 6-7 days pc, a higher severity was observed in animals from the FIV group (combined total score 24), compared to the Ad-GFP control (combined total score 10). This included eosinophilic infiltrate and perivascular cuffing that was not observed in the control-vaccinated ferrets. The lung pathology in the ferret model^24^ was quite transient and thus at 13-15 days pc, whilst there was some individual variability present, both groups showed mild pathology.

Multifocal mild to moderate hepatitis has also been described as a potential adverse effect of SARS-CoV-1 vaccines^45^. However, these lesions are found as a background finding for this species in many experimental studies, although viral infections, systemic or in the gastrointestinal tract, have also been related to the presence of these periportal inflammatory infiltrates^46^. Due to the variability in severity and the fact that naïve ferrets frequently show some degree of hepatitis, the interpretation of this lesion must be taken cautiously.

Following this observation of mild transient enhanced disease in the two FIV-vaccinated ferrets culled at day 7, we have tested the same FIV in 6 macaques along with 6 unvaccinated controls. We have previously compared the course of SARS-CoV-2 infection in both rhesus and cynomolgus macaques and showed virus replication in the upper and lower respiratory tract with pulmonary lesions resembling mild COVID-19 in humans^25^. Macaques allow for a more detailed examination of lung pathology using precise scoring system devised in our recent study. We also have considerable experience of in life CT scanning of macaques^47 30 25^ which allows further aspects of lung pathology to be characterised at time points before necropsy. This detailed analysis has revealed no evidence of enhanced disease in macaques at any time point, but rather that the FIV provided some protection against the mild-to-moderate lung pathology observed in the unvaccinated control macaques.

The presence of inflammatory infiltrates, and particularly perivascular cuffing, has been described as a feature potentially related to VED in SARS-CoV-1 preclinical vaccine trials^19 21 22^. In our study these infiltrates were always of mild to moderate severity.

Characterisation of the immune response to the FIV vaccine prior to challenge in both species confirmed the expectation of modest immunity to SARS-CoV-2 spike. A significant rise in anti-S IgG, but not anti-N IgG was detected by ELISA. The geometric mean neutralising titre of 89 seen in ferrets and 61 in macaques was low compared to that observed in primate studies with candidate vaccines^10 8 7 11^ and in clinical trials^29 7 48 4 5^. However, the larger rise in neutralising antibody titre in FIV-vaccinated ferrets and macaques following challenge, compared to control animals indicated that priming had been mediated by the FIV. The SARS-CoV-2-specific interferon γ response measured in splenocytes using an ELISpot assay 6 or 7 days after challenge showed a greater response in the FIV-vaccinated animals to live virus, membrane and nucleocapsid peptide pools but very little response to the S protein pools, indicating a poor cellular response to the S antigen with FIV. Conversely, spike peptide-specific IFNγ SFU measured in macaques increased significantly following FIV vaccination and evaluation of the spike-specific interferon-γ response in splenocytes and PBMC collected from macaques at a similar time after SARS-CoV2 challenge revealed the reverse pattern to that observed in ferrets, with greater responses measured in the unvaccinated group relative to the FIV-vaccinated animals. This may reflect that improved priming of the T-cell mediated response contributed to the protection afforded by the FIV vaccine in this species if response magnitude is driven by antigenic load following infection.

The expression of checkpoint inhibitory receptors is often considered a marker of T-cell exhaustion, although more recently PD-1 signaling has also been linked to improved effector T-cell priming and enhanced clearance of acute viral infections^49^. Similarly, the induction of regulatory CD4 T-cells, and the inhibitory influence they are likely to exert, may be considered counterproductive to vaccine induced immunity. However, the increased frequency of T-regs observed following FIV vaccination is likely to reflect the immune response typically induced by coronaviruses in this species, as similar increases were also seen in the unvaccinated animals early after SARS-CoV2 infection, and thus may help to explain the relatively mild disease that develops in this species and the improved outcome observed in FIV vaccine primed macaques^50 51^.

There are limitations to the current study, the principal of which is the small number of ferrets in the early culled group. The transient nature of the pathology meant that these differences resolved by the second necropsy time point.

Some insight into the weak anti-S neutralising response induced by the FIV was gained using a capture ELISA which preserved the conformation of the S trimer on the solid phase, unlike direct coating onto the ELISA plate which modified antigenicity (data not shown), as has been observed for soluble forms of the HIV-1 envelope glycoprotein trimer^52^. Formaldehyde treatment of S trimer in this format was the same as that used to inactivate the vaccine, allowing extrapolation of ligand binding to the ELISA-captured formaldehyde-treated S trimer to that on the virus.

Formaldehyde cross-linking resulted in a 2-fold reduction in binding of ACE2-Fc and two RBD-specific MAbs (CR3022 and EY6A) to formaldehyde-treated compared to untreated S trimer. By contrast, formaldehyde treatment of recombinant RBD did not affect binding of ACE2-Fc or these MAbs, implying that formaldehyde treatment cross-linked a proportion of S trimer into a non-ligand binding conformation. These results are consistent with the location of lysine and arginine residues, and suggest that RBD exposure may be limited by cross-linking, reducing exposure of neutralising antibody epitopes on the RBD. This result may be of more general interest, since other viral envelope glycoproteins, such as those of HIV-1, are metastable and sample different conformational states, some of which are more relevant to neutralisation than others^53^. Cross-linking may trap these different conformational states, modifying exposure of neutralising antibody epitopes to B cell recognition^54^.

Formaldehyde inactivation has been widely used to prepare inactivated viral vaccines^55^ and as a toxoiding agent for bacterial toxin vaccines. Some of the considerations for inactivated SARS-CoV-2 vaccines are discussed in a recent commentary^56^. The study reported here has confirmed that caution should be used if formaldehyde is the inactivation reagent for COVID-19 vaccines. Several vaccines are in development that use β-propiolactone as inactivation agent. One such vaccine has been shown to be protective in rhesus macaques following SARS-CoV-2 challenge without induction of VED^7^. A preliminary report of phase 1 and 2 studies with another β-propiolactone vaccine indicated that it is tolerated, safe and produced neutralising antibodies in phase 1 and 2 studies^48^. In addition, the authors mention in the discussion that enhancement of disease was not observed in primates following SARS-CoV-2 challenge but no pathology results are presented.

In conclusion, we have prepared an experimental SARS-CoV-2 vaccine based on previous inactivated virus studies that induced VED. We showed no evidence of enhanced disease at later time points in ferrets or at any time in a more in-depth analysis in rhesus macaques which included CT imaging. However, we did observe increased pathology scores, early in the infection of FIV-vaccinated ferrets which resolved by the later necropsy time point. It is reassuring that, even with a vaccine deliberately designed to induce enhanced disease, no enhancement was seen apart from at 7 days post infection in ferrets. Future studies to investigate the potential of SARS-CoV-2 vaccines to cause enhanced disease should examine lung pathology at multiple time points including soon after challenge. Formalin-inactivated virus can be used as a suboptimal comparator to determine the potential of SARS-CoV-2 vaccine candidates to induce VED so that unsuitable vaccines are identified at an early stage of development before significant clinical studies commence.

## Acknowledgements

The authors acknowledge the contributions of all staff within the PHE Biological Investigations Group for assistance with the delivery of the *in vivo* study and B Cavell, J Gouriet, V Lucas, D Ngabo, S Thomas and R Watson for assistance with processing of *in vivo* samples. The authors thank S Findlay-Wilson, T Hender, N McLeod and C Turner for their assistance with RNA extraction and PCR, E Penn for assistance with neutralisation assays and T Tipton for preparation of ELISpot peptides. Alexandra Morrison, Adam Mabbut and Dinos Gkolfinos assisted with ELISPot assays. The authors thank M Elmore and M Matheson for assistance with data analysis. The authors also acknowledge the kind donation of the S trimer expression construct by R. Shattock and P. McCay, the ACE2-Fc construct by H. Waldmann, and the RBD-specific monoclonal antibodies CR3022 and EY6A by T. Tan and K-Y. Huang respectively. We thank T. Schiffner for assistance with the molecular modelling and S. Zhang for assistance with S trimer production and the Jenner Institute Viral Vector Core Facility for the production of Ad-GFP. Q. Sattentau, S. Gilbert and T. Lambe are Jenner Institute Investigators. Q. Sattentau is a James Martin Senior Fellow. The work was supported by UKRI Grant MC_PC_19080 and MRC UKRI Grant MC_PC_19055. Viral stock preparation was funded by the Coalition for Epidemic Preparedness Innovations.

## Author contributions

Kevin R. Bewley, development of virus growth, plaque and neutralisation assay methods. Design of virus inactivation strategy, vaccine preparation, drafting manuscript and data analysis; Karen Gooch, study plan and management, data analysis and manuscript review; Kelly M. Thomas, study plan and management, vaccine preparation, data collation and analysis, PBMC preparation, manuscript review; Stephanie Longet, ELISA design, assay and analysis; Nathan Wiblin, in vivo study supervisor; Laura Hunter, histopathology slide preparation and staining; Kin Chan, SDS-PAGE and western blots; Phillip Brown, ELISpot assay and analysis, processing of in vivo samples; Rebecca A. Russell, performed experimental procedures, analysis and preparation of figures; Catherine Ho, Preparation of in vivo samples and sera; Gillian Slack, PCR data analysis; Holly E. Humphries, PRNT assays and analysis; Leonie Alden, in vivo study management; Lauren Allen, performed PRNT assays; Marilyn Aram, performed PCR assays; Natalie Baker, virus inactivation and confirmation; Emily Brunt, performed PRNT assays; Rebecca Cobb, processing of in vivo samples; Susan Fotheringham, in vivo study management; Debbie Harris, in vivo study management; Chelsea Kennard, histology analysis including RNAscope; Stephanie Leung, performed PRNT assay; Kathryn Ryan, processed in vivo samples; Howard Tolley, performed electron microscopy; Nadina Wand, RNA extraction and PCR assays; Andrew White, Laura Sibley and Charlotte Sarfas designed and performed the macaques ELISpot assays; Xiaochao Xue, prepared and characterised proteins, manuscript drafting; Yper Hall, preparation and characterisation of inactivated virus; Teresa Lambe, in vivo study design and provision of control vaccine; Sue Charlton, in vivo study design and preparation of inactivated vaccine; Simon Funnell, study design and manuscript preparation; Sarah Gilbert, in vivo study design and provision of control vaccine; Quentin J. Sattentau, designed and performed experiments to characterise the effect of formaldehyde on SARS-CoV-2 spike protein, drafted manuscript; Francisco J. Salguero, lead the histopathology analysis and reporting, drafted manuscript; Geoff Pearson, histopathology analysis; Emma Rayner, histopathology analysis; Sally Sharpe, CT scanning and analysis, manuscript preparation; Fergus Gleeson, analysis of CT scans; Andrew Gorringe, scientific direction and lead preparation of the manuscript; Miles Carroll, established the project, in vivo study design, scientific direction and manuscript preparation.

## Data availability statement

The data that support the findings of this study are available from the corresponding authors upon reasonable request.

