## Supplemental figures 1-4 for "Immunological and pathological outcomes of SARS-CoV-2 challenge after formalin-inactivated vaccine immunisation of ferrets and rhesus macaques"

Supplementary figure 1

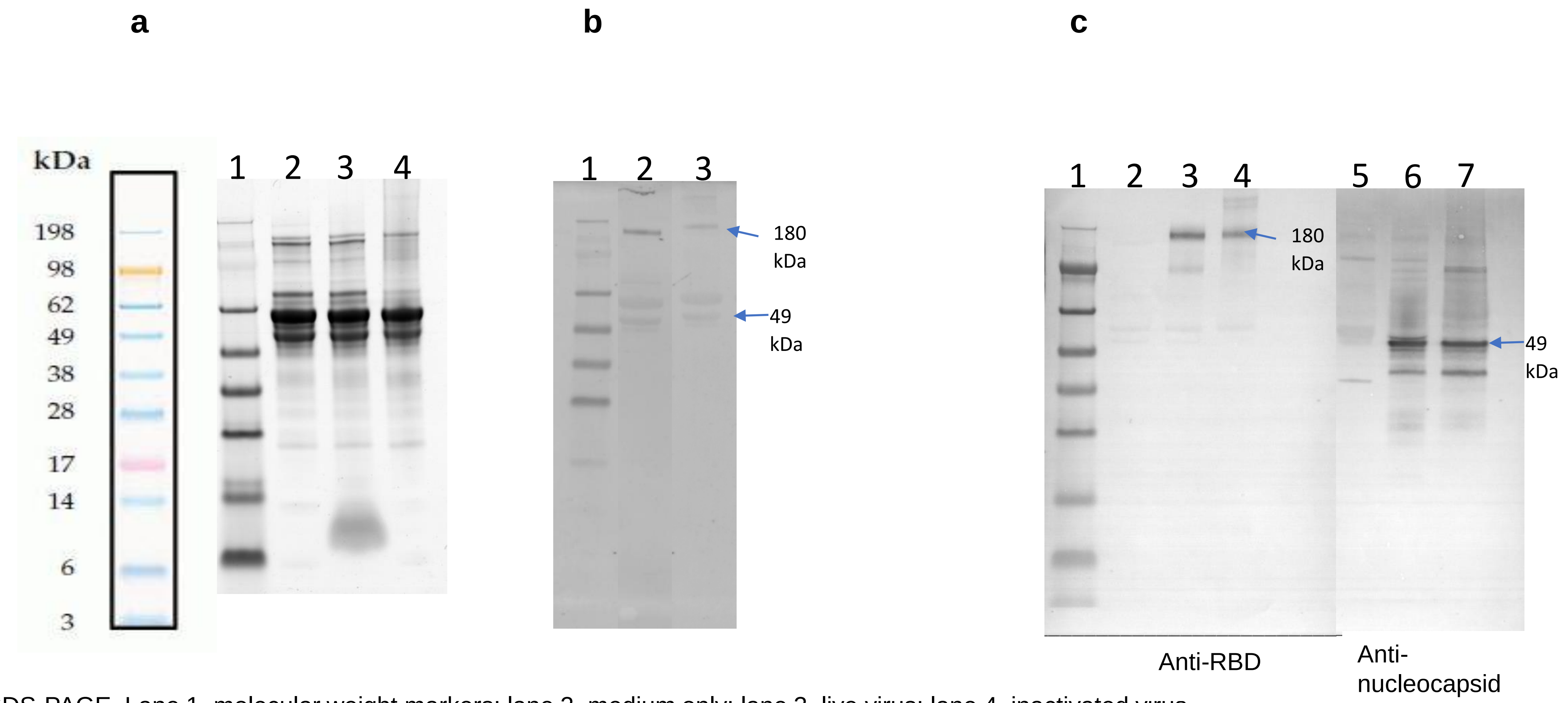


a. SDS-PAGE. Lane 1, molecular weight markers; lane 2, medium only; lane 3, live virus; lane 4, inactivated virus.
b. Western blot probed with MERS convalescent S3 serum. Lane 1, molecular weight markers; lane 2, live virus; lane 3 inactivated virus.
c. Western blot probed with anti-RBD (lanes 2-4) or anti-nucleocapsid polyclonal antiserum (lanes 5-7). Lane 1, molecular weight markers; lane 2, medium only; lane 3, live virus; lane 4, inactivated virus; lane 5, medium only; lane 6, live virus; lane 7 inactivated virus.

Supplementary figure 2

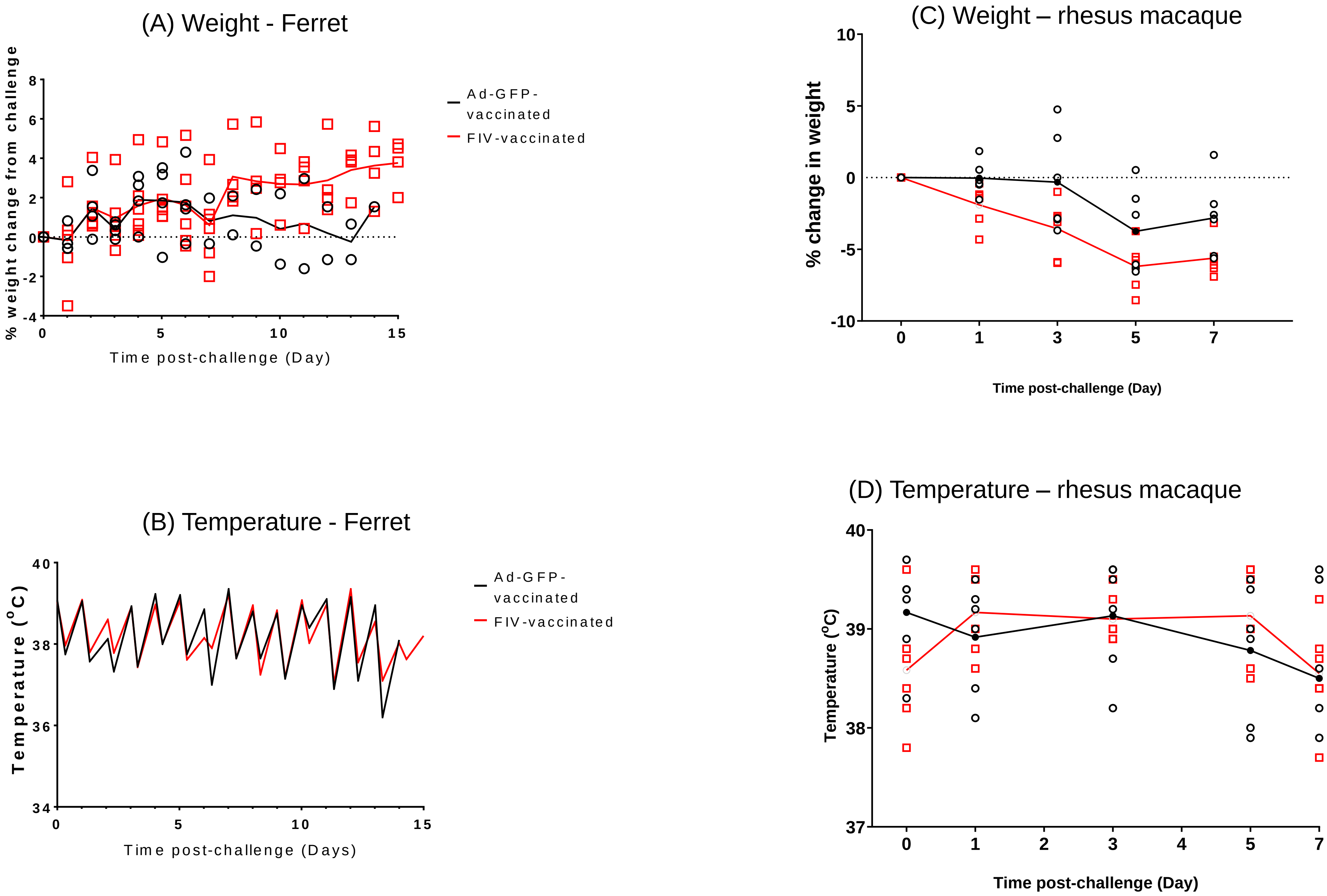

Supplementary figure 3

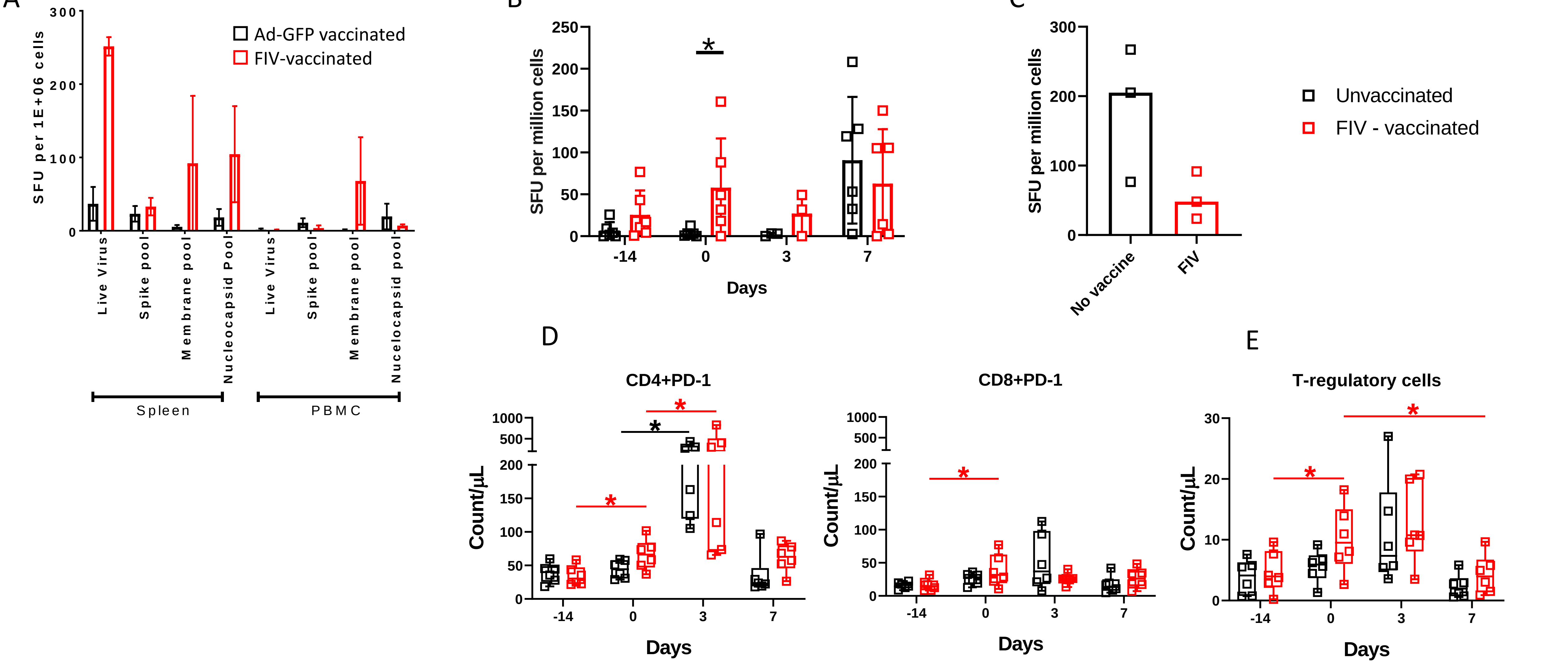


SARS-CoV-2-specific IFNγ spot forming unit (SFU) frequency measured by ELISPOT in PBMCs and splenocytes following vaccination with formaldehyde-inactivated virus (FIV) or control/no vaccine and subsequent infection with SARS-CoV2 . A) SARS-CoV-2-specific SFU in splenocytes and PBMCs isolated from ferrets 6-8 days following SARS-CoV2 infection. B) Spike-specific SFU in PBMCs isolated from macaques. FIV vaccination administered at timepoint Day -14, SARS-CoV-2 infection at Day 0. C) Spike-specific SFU measured in splenocytes 6-8 days following SARS-CoV-2 infection. Black = no vaccine, Red = FIV. Each square represents one animal and bar at median. Mann-Whitney U-test significance values shown ($p \leq 0.05$). D) Quantification of $CD4^{+}$ and $CD8^{+}$ T cells expressing PD-1 prior to (day -14), 14 days after FIV vaccination (day 0) and at days 3 and 6-8 days (day 7) following SARS-CoV2 infection in NHPs. E) Quantification of NHP CD4+ T-regulatory cells determined by expression of CD25 and CD127 by whole blood immunophenotyping flow cytometry assay. Box plots show the group median +/- the inter-quartile range, with minimum and maximum values connected by whiskers. Asterisks denote significant differences determined by Wilcoxon signed rank test for paired comparisons, (*) $p \leq 0.05$.

### Supplementary Figure 4

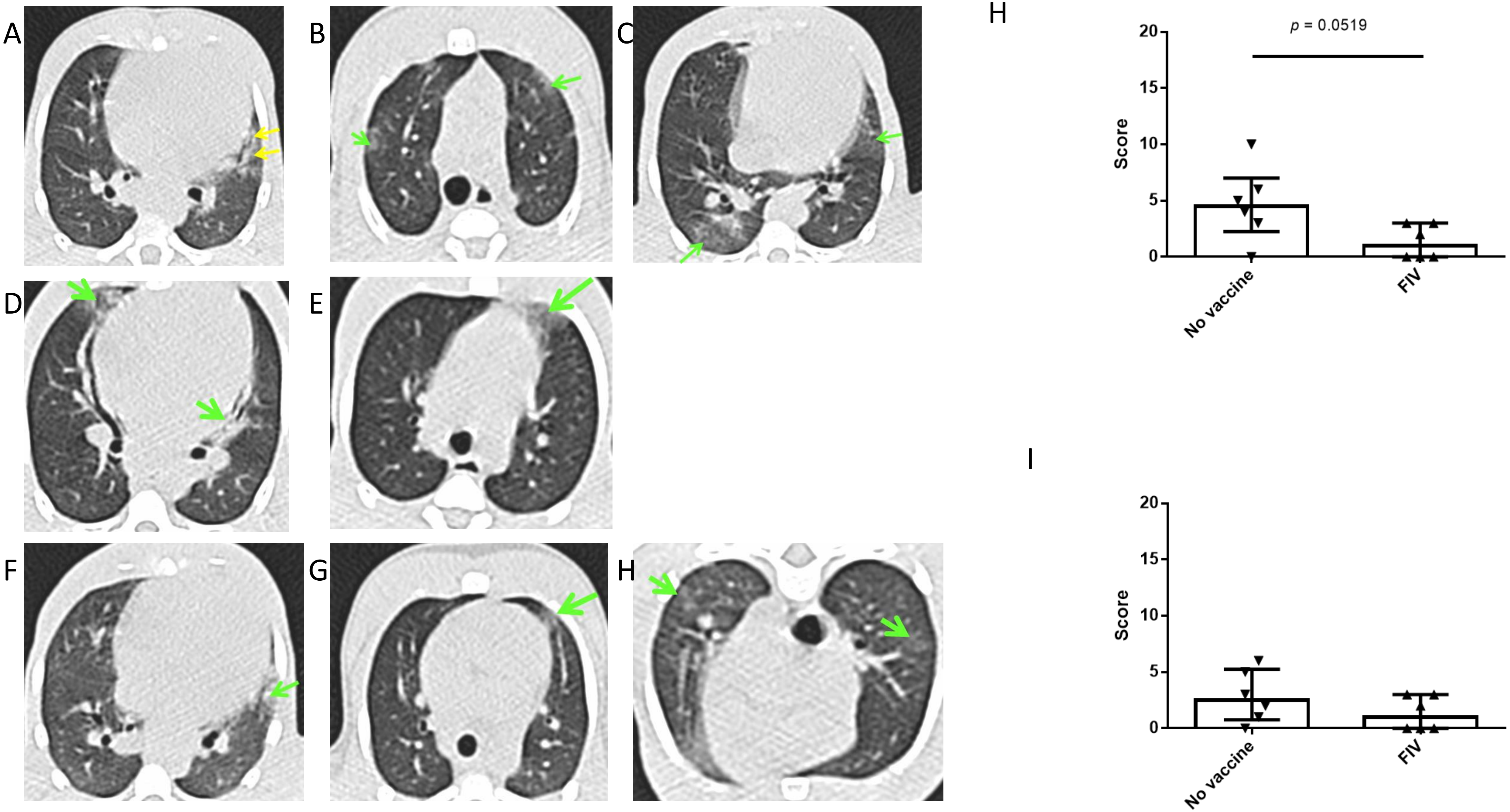


Representative example of pulmonary abnormalities identified on images constructed from CT scans five days after SARS-CoV-2 challenge collected from macaques that did not receive a vaccination A – E and macaques that received FIV vaccine F, G, H. Green arrows indicate areas of ground glass opacification, yellow arrows indicate areas of consolidation. Images from macaques that did not have abnormal features are not shown. Plots show the scores for attributed for possession of characteristic features associate with COVID (pattern score) [H] and distribution of the abnormalities through the lung (zone score) [I] in non vaccinated and FIV-vaccinated macaques showing non-significant trends for reduction in severity in the FIV group (pattern score $p = 0.0519$, zone score $p = 0.3052$) Mann-Whitney U-tests; Box plots show the experimental group median with +/- IQR indicated by box whiskers, symbols show scores measured in individual animals.
